# Epigenetic control of microglial mitochondrial immunity by KAT7 drives Alzheimer’s disease pathogenesis

**DOI:** 10.64898/2026.02.19.706884

**Authors:** Yongqing Liu, Minghua Fan, Yingzhi Ye, Henry Yi Cheng, Shuying Sun, Zhaozhu Qiu

## Abstract

Mitochondrial DNA (mtDNA)-driven innate immune signaling sustains chronic neuroinflammation in neurological diseases such as Alzheimer’s disease (AD), yet how this pathway is regulated in microglia remains poorly understood. Here, we identify the histone acetyltransferase KAT7 (HBO1) as a central epigenetic regulator that links chromatin remodeling to mitochondrial immune activation. KAT7 and its histone mark H3K14ac are elevated in microglia from 5×FAD mice and human AD brains. Integrative transcriptomic and epigenomic analyses reveal that KAT7 activates transcription of *Cmpk2*, a mitochondrial kinase essential for mtDNA synthesis. Loss of KAT7 reduces *Cmpk2* expression, impairs mtDNA replication and release, and consequently suppresses cGAS-STING and NLRP3 signaling. Importantly, both microglia-specific deletion and pharmacological inhibition of KAT7 mitigate cytosolic mtDNA-induced neuroinflammation, decrease amyloid-β burden, restore synaptic plasticity, and improve cognitive function in 5×FAD mice. Together, these findings uncover an epigenetic-mitochondrial axis sustaining microglial pathogenicity and establish KAT7 as a promising therapeutic target for AD.

## INTRODUCTION

Alzheimer’s disease (AD) is the most common cause of dementia, affecting tens of millions worldwide and imposing an ever-growing societal and economic burden^1^. It is neuropathologically characterized by the accumulation of extracellular amyloid-β (Aβ) plaques and intracellular neurofibrillary tangles of hyperphosphorylated tau^2^. Yet, decades of research targeting these hallmark proteins have yielded only modest clinical benefit, indicating that additional mechanisms underlie AD pathogenesis^3^. Mounting evidence implicates chronic neuroinflammation as a central disease mechanism^4-6^. Genome-wide association studies have linked many AD risk loci to microglial pathways, highlighting microglia, the brain-resident immune cells, as key players in AD^7-9^. While microglia initially protect the brain by clearing misfolded proteins, chronic activation drives their transition into a proinflammatory state marked by excessive cytokine production and loss of homeostatic functions^10^. This maladaptive state fuels a self-perpetuating cycle of inflammation and neurodegeneration.

Mitochondrial dysfunction is increasingly recognized as a critical driver of chronic microglial inflammation^11-13^. In both aging and AD brains, microglia exhibit elevated levels of mitochondrial DNA (mtDNA) in the cytosol, where it acts as a potent damage-associated molecular pattern^14-16^. Cytosolic mtDNA is sensed by cyclic GMP-AMP synthase (cGAS), which produces the second messenger cyclic GMP-AMP (cGAMP) to activate STING (stimulator of interferon genes)^12^. Activation of this pathway promotes phosphorylation of TBK1 (TANK-binding kinase 1) and IRF3 (interferon regulatory factor 3), leading to type I interferon induction and the release of proinflammatory cytokines^17^. Persistent cGAS-STING activation in microglia sustains a maladaptive inflammatory state that drives AD progression. Notably, genetic or pharmacological inhibition of this signaling pathway mitigates microglial activation and alleviates AD-related pathology^15,18^, underscoring its pathogenic role. However, the upstream mechanisms governing this cytosolic mtDNA-initiated inflammatory cascade in microglia remain largely unknown.

Epigenetic regulation provides a critical layer of control over gene expression, enabling transient stimuli to be converted into long-lasting transcriptional programs^19-21^. Among these mechanisms, histone acetylation plays a pivotal role: acetylation of lysine residues on histone tails generally relaxes chromatin structure and facilitates transcription^22^. This places histone acetyltransferases (HATs), also known as lysine acetyltransferases (KATs), at the core of transcriptional reprogramming^22^; however, their contribution to microglial inflammation and AD pathogenesis remains poorly understood. Here, through gene expression profiling, we identify KAT7 (also known as HBO1), a member of the MYST family of HATs^23^, as a key epigenetic regulator of neuroinflammation. By coupling histone acetylation to enhanced mitochondrial DNA synthesis and cGAS-STING activation in microglia, our work uncovers a previously unrecognized epigenetic mechanism driving chronic neuroinflammation and highlights KAT7 as a promising therapeutic target for AD.

## RESULTS

### Expression of the KAT7 complex is upregulated in microglia from both 5×FAD mouse model and human AD brains

To determine whether epigenetic regulators of histone acetylation are altered during microglial activation, we analyzed public RNA-seq data from primary mouse microglia^24^ and found that *Jade2*, which encodes a scaffold subunit of the KAT7 complex, was selectively upregulated in response to lipopolysaccharide (LPS) stimulation (**Fig. 1A**). We next performed RNA-seq on LPS-treated BV2 cells, a mouse microglia-derived cell line, and also observed increased expression of *Jade2* in response to LPS (**Fig. 1B**), whereas expression of other HAT complexes remained largely unchanged. These findings were further validated by qPCR and western blot analyses (**Fig. 1C** and **Suppl. Fig. 1**). Given that neuroinflammation is a hallmark of AD^5^, we examined whether expression and activity of the KAT7 complex are also elevated in this context. To this end, we isolated microglia from 6-month-old 5×FAD mice, a well-characterized AD model carrying five familial AD mutations^25^, using CD11b microbeads (**Suppl. Fig. 2A-B**). qPCR analysis revealed that the expression of *Kat7* and two of its scaffold subunits, *Jade2* and *Brpf2*, was increased in microglia from 5×FAD mice compared with age-matched wild-type (WT) littermate controls (**Fig. 1D**). RNAscope analysis further showed upregulation of *Kat7* and *Jade2* specifically in microglia, but not in neurons or astrocytes (**Suppl. Fig. 2C-D**). To extend these findings to humans, we analyzed a publicly available RNA-seq dataset (GSE125050) from post-mortem superior frontal gyrus tissue of AD patients and healthy controls^26^. While expression of *KAT7* and its other subunits was unchanged, the scaffolds *JADE2* and *BRPF3* were upregulated in microglia from AD patients (**Suppl. Fig. 2E**). Because suitable antibodies for KAT7 complex proteins were unavailable, we leveraged the fact that KAT7 is the primary enzyme responsible for H3K14 acetylation (H3K14ac) in cells^27-29^ and performed immunostaining with an anti-H3K14ac antibody on brain sections. Consistently, whereas microglial H3K14ac levels were low in WT controls, they were markedly elevated in 5×FAD mice (**Fig. 1E**), with no changes observed in neurons or astrocytes (**Suppl. Fig. 3**). Importantly, H3K14ac signals were also strongly increased in microglia from postmortem human AD brains compared with healthy controls (**Fig. 1F**). Together, these results demonstrate that upregulation of the KAT7 complex and its histone acetylation mark accompanies microglial activation, implicating KAT7 in neuroinflammation and AD pathogenesis.

**Fig. 1.**
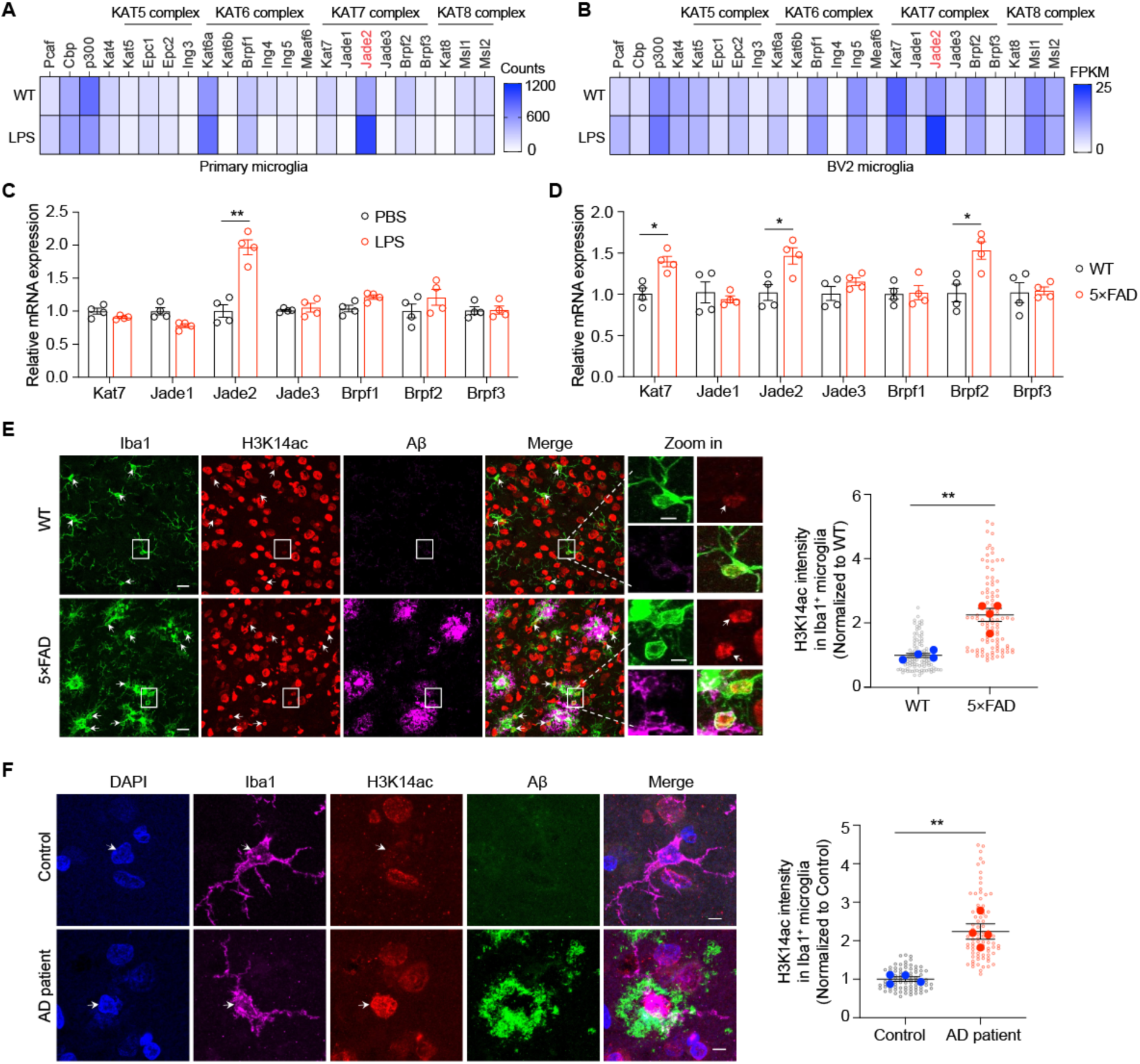
Expression of the KAT7 complex is elevated in microglia from both 5×FAD mouse model and human AD patients. **A**, Heatmap of HATs components from public RNA-seq dataset (GSE90046, n=3 per group) in mouse primary microglia treated with LPS. **B**, Heatmap of HATs components from RNA-seq data (n=3 per group) in BV2 microglia treated with LPS. **C**, qPCR analysis of KAT7 complex in BV2 cells treated with LPS. n=4 per group. Unpaired student’s t test. **D**, qPCR analysis of KAT7 complex in the isolated microglia from 6-month-old 5×FAD mice. n=4 per group. Unpaired student’s t test. **E**, Left: Representative images of H3K14ac co-stained with Aβ plaques and microglia (Iba1) in the cortex region of 6-month-old WT and 5×FAD mice. Scale bar, 20 µm (left), 5 µm (right). Right: Quantification of H3K14ac intensity in microglia. n=100 cells from 4 mice per group. Mann-Whitney test. **F**, Left: Representative images of H3K14ac co-stained with Aβ plaques and microglia in the frontal lobe region of AD patients and healthy controls. Scale bar, 6 µm. Right: Quantification of H3K14ac intensity in microglia. n=80 cells from 4 samples per group. Mann-Whitney test. White arrowheads indicate H3K14ac in microglia. *p<0.05, **p<0.01, ***p<0.001. Data are presented as mean ± SEM.

### KAT7 regulates LPS- and Aβ-induced inflammatory responses in microglia

To investigate the role of KAT7 in neuroinflammation, we generated *Kat7*-knockout (KO) BV2 microglial cells using CRIPSR-Cas9 (**Fig. 2A**) and employed LPS stimulation as a well-established model of inflammatory activation. *Kat7* deletion markedly reduced LPS-induced iNOS expression and secretion of the inflammatory cytokine IL-6 (**Fig. 2B-D**). Conversely, overexpression of WT KAT7 enhanced IL-6 production, whereas the catalytically inactive mutant (KAT7-E508Q) failed to do so (**Fig. 2E-F**), indicating that the enzymatic activity of KAT7 is required for its pro-inflammatory function. We next examined the role of the scaffold subunit JADE2 in neuroinflammation. Overexpression of JADE2 increased KAT7 protein levels, suggesting that JADE2 stabilizes the KAT7 complex (**Suppl. Fig. 4A**). Moreover, JADE2 promoted the expression of pro-inflammatory factors in a manner dependent on its interaction with KAT7 (**Suppl. Fig. 4B**). To validate these findings in primary cells, we cultured microglia from neonatal mice and transfected them with *Kat7*-specific siRNAs (**Fig. 2G**). Consistently, *Kat7* knockdown significantly attenuated LPS-induced IL-6 production (**Fig. 2H-J**). Similarly, pharmacological inhibition of KAT7 with WM-3835, a potent small-molecule inhibitor^27^, suppressed LPS-induced *Nos2* and *Il6* expression in a dose-dependent manner (**Fig. 2K**). In addition to LPS, we further asked whether KAT7 regulates inflammation driven by aggregated Aβ. Treatment of primary mouse microglia with oligomeric Aβ42 robustly induced production of the pro-inflammatory cytokines IL-6 and IL-1β, which was markedly attenuated by WM-3835 (**Fig. 2L-O**), supporting a role of KAT7 in mediating Aβ-induced inflammatory responses. These results establish KAT7 as a critical driver of microglial inflammatory responses to both LPS and aggregated Aβ.

**Fig. 2.**
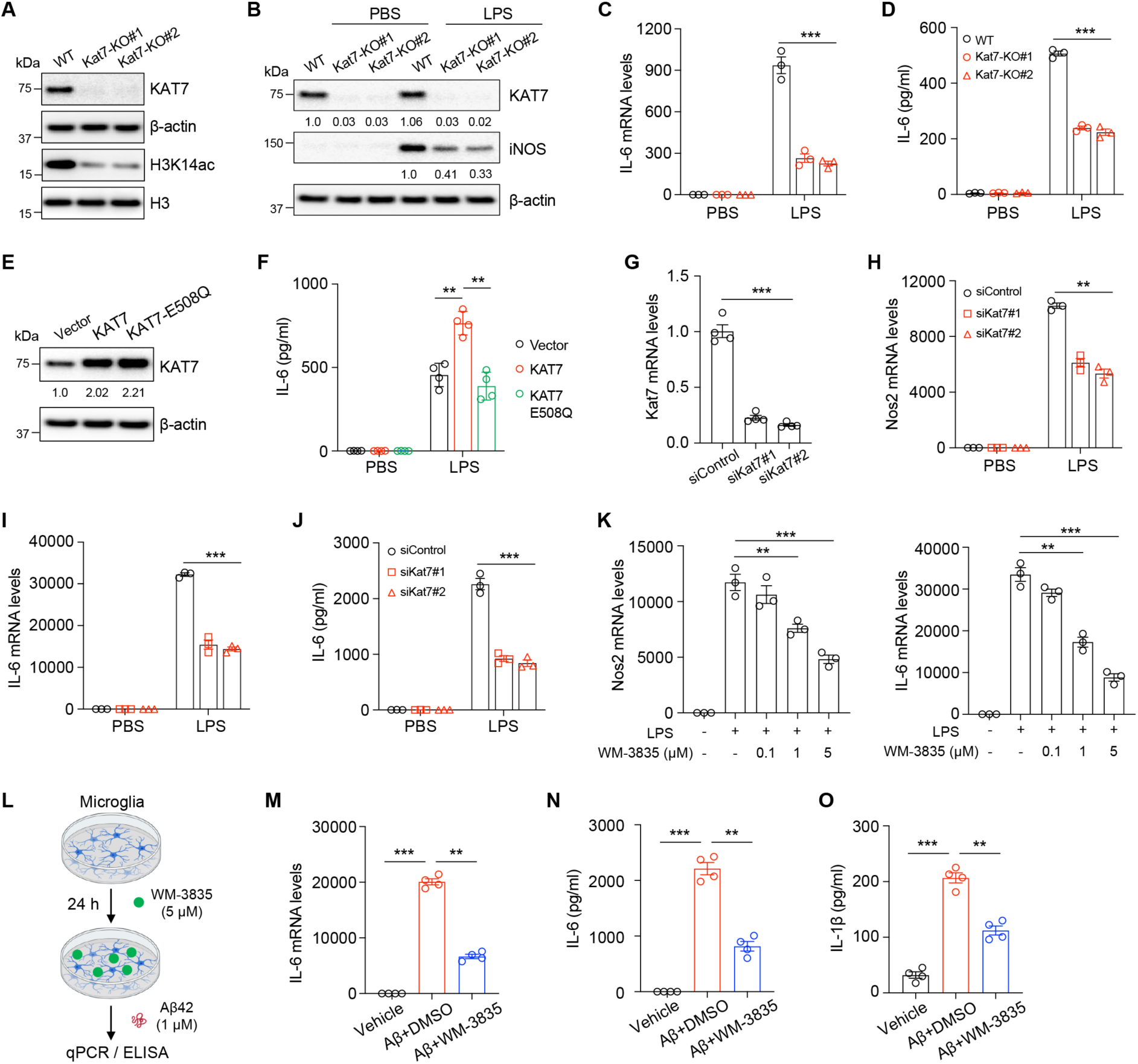
KAT7 regulates LPS- and Aβ- induced inflammatory responses in microglia. **A**, Generation of *Kat7*-KO BV2 cells. **B**, Western blot analysis of KAT7 and iNOS levels in *Kat7*-KO BV2 cells treated with LPS. **C-D**, qPCR and ELISA analysis of IL-6 levels in *Kat7*-KO BV2 cells treated with LPS. n=3. **E**, Western blot analysis of KAT7 overexpression in BV2 cells. **F**, ELISA analysis of IL-6 secretion in BV2 cells treated with LPS. n=4. **G**, qPCR analysis of *Kat7* level in mouse primary microglia transfected with siRNA. n=4. **H-I**, qPCR analysis of *Nos2* and *Il-6* levels in primary microglia treated with siRNA and LPS. n=3. **J**, ELISA analysis of IL-6 secretion in primary microglia. n=3. **K**, qPCR analysis of *Nos2* and *Il-6* levels in primary microglia treated with WM-3835 and LPS. n=3. **L**, Schematic diagram showing mouse primary microglia treated with WM-3835 and oligomeric Aβ42. **M**, qPCR analysis of *Il-6* levels in primary microglia treated with WM-3835 and oligomeric Aβ42. n=4. **N-O**, ELISA analysis of IL-6 secretion (6 h after Aβ42 treatment) and IL-1β (24 h after Aβ42 treatment) in primary microglia treated with WM-3835 and Aβ42. n=4. **p<0.01, ***p<0.001. One-way ANOVA test (**G**, **K**, and **M**-**O**). Two-way ANOVA test (**C**, **D**, **F**, and **H**-**J**). Data are mean±SEM.

### Combined transcriptomic and epigenomic profiling reveals *CMPK2* as a critical transcriptional target of KAT7

To elucidate the molecular mechanisms by which KAT7 regulates neuroinflammation, we performed RNA-seq to profile transcriptomic changes in WT and *Kat7* KO BV2 cells with or without LPS treatment (**Fig. 3A**). Using an adjusted p-value < 0.05 and |log_2_ fold change| > 1 as cutoffs, we identified 1,074 upregulated genes in response to LPS stimulation in WT cells (**Fig. 3B**). Deletion of *Kat7* attenuated the induction of 110 of these genes (**Fig. 3C-D**). Gene ontology (GO) analysis revealed that these KAT7-dependent genes were significantly enriched in pathways related to interferon signaling and inflammatory responses (**Fig. 3D**). In contrast, of the 276 genes downregulated by LPS in WT cells, only five were reversed by *Kat7* deletion (**Suppl. Fig. 5A**), indicating that KAT7 has minimal impact on LPS-induced transcriptional repression. Given that KAT7 is primarily responsible for H3K14 acetylation, a histone mark associated with transcriptional activation^29^, we next performed CUT&Tag (Cleavage Under Targets and Tagmentation) analysis to map the genome-wide distribution of H3K14ac (**Fig. 3E**). Genomic distribution analysis revealed that most differential H3K14ac peaks were located in promoter regions: 38% (6,121 peaks) between WT_LPS and WT cells, and 40% (5,776 peaks) between KO_LPS and WT_LPS cells (**Suppl. Fig. 5B-C**). Among the genes showing increased H3K14ac enrichment at promoters in WT cells upon LPS stimulation, 244 exhibited reduced acetylation in *Kat7* KO cells (**Fig. 3E** and **Suppl. Fig. 5D**). Integrative analysis of RNA-seq and CUT&Tag datasets identified 17 genes as potential direct transcriptional targets of KAT7 in response to LPS stimulation (**Fig. 3F-G**). The limited overlap likely reflects the stringent thresholds applied. Validation by qPCR and quantitative chromatin immunoprecipitation (qChIP) using KAT7 and H3K14ac antibodies (with H3K23ac antibody as a negative control) confirmed the profiling results (**Fig. 3H-I** and **Suppl. Fig. 5E-G**). Notably, among the KAT7-regulated genes, *Cmpk2* (Cytidine/uridine monophosphate kinase 2) was one of the most dramatically suppressed in *Kat7* KO cells in response to LPS, highlighting it as a key downstream target (**Fig. 3H** and **Suppl. Fig. 5E**). Importantly, *Cmpk2* was also robustly induced by oligomeric Aβ42 in primary mouse microglia, and this induction was markedly attenuated by *Kat7* knockdown (**Fig. 3J**). Collectively, these findings identify *Cmpk2* as a direct transcriptional target of KAT7 and suggest that it functions as a shared downstream effector in microglial inflammatory responses to both LPS and Aβ stimulation.

**Fig. 3.**
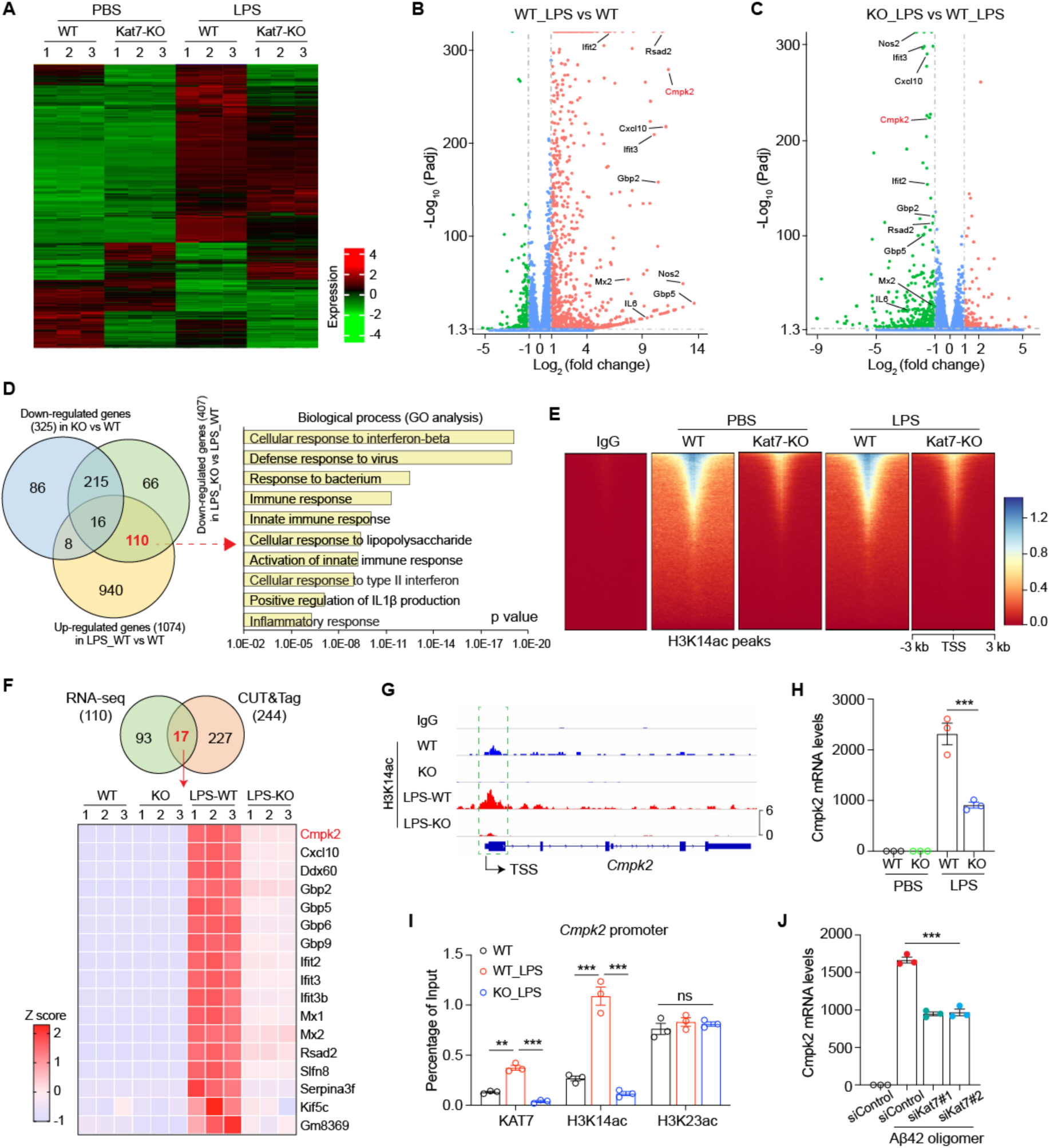
Integrated transcriptomic and epigenomic analysis identifies *Cmpk2* as a key target of KAT7. **A**, Heatmap of RNA-seq in WT and *Kat7*-KO BV2 cells with or without LPS stimulation. n=3 replicates per group. **B**, Volcano plot showing differentially expressed genes between WT_LPS and WT BV2 cells. **C**, Volcano plot showing differentially expressed genes between *Kat7*-KO_LPS and WT_LPS BV2 cells. **D**, Venn diagram (left) of overlapping genes among downregulated in KO vs WT, downregulated in KO_LPS vs WT_LPS, and upregulated in WT_LPS vs WT. Right: GO pathway analysis of the 110 overlapped genes. **E**, Heatmap of CUT&Tag with IgG and H3K14ac antibodies in WT and *Kat7*-KO BV2 cells treated with or without LPS stimulation. **F**, Top: Venn diagram showing overlapped genes of RNA-seq (110 genes) and CUT&Tag (244 genes). Bottom: Heatmap of 17 overlapped genes in RNA-seq. **G**, Representative CUT&Tag tracks of H3K14ac in *Cmpk2*. Green box indicated proximal promoter. TSS, transcriptional start site. **H**, qPCR analysis of *Cmpk2* in BV2 cells treated with LPS. n=3. **I**, qChIP analysis of *Cmpk2* promoter using the indicated antibodies in BV2 cells. n=3. **J**, qPCR analysis of *Cmpk2* in primary microglia treated with Kat7 siRNA and oligomeric Aβ42. n=3. **p<0.01, ***p<0.001. One-way ANOVA test. Data are mean±SEM.

### KAT7 drives microglial innate immune signaling via CMPK2-dependent mtDNA synthesis

*CMPK2* encodes a mitochondrial nucleotide monophosphate kinase that acts as the critical rate-limiting enzyme, ensuring dNTP precursor availability and driving the dramatic upregulation of mitochondrial DNA (mtDNA) synthesis during macrophage activation^30^. CMPK2-dependent mtDNA synthesis facilitates the release of mtDNA into the cytoplasm, which subsequently activate the cGAS-STING signaling pathway and the NLRP3 inflammasome^30-32^. Given that KAT7 regulates *Cmpk2* expression during microglia activation, we tested whether it modulates mtDNA replication, release, and downstream innate immune responses. We first performed 5-ethynyl-2’-deoxyuridine (EdU) labeling, which preferentially incorporates into newly synthesized mtDNA and appears as bright cytoplasmic puncta in non-proliferating primary mouse microglia (**Fig. 4A-B**). *Kat7* knockdown impaired LPS-induced mtDNA replication, an effect rescued by WT CMPK2 but not by a catalytically inactive CMPK2 mutant (CMPK2-D330A) (**Fig. 4A-B** and **Suppl. Fig. 6**), indicating that KAT7 promotes CMPK2-dependent mtDNA synthesis in response to LPS. To assess mtDNA release, we primed microglia with LPS followed by ATP treatment, a common method to induce mitochondrial stress and trigger innate immunity^32^. qPCR analysis of cytosolic fractions using primers specific to the mitochondrial D-loop region revealed elevated mtDNA levels in control microglia (**Fig. 4C**), indicating increased release from damaged mitochondria. Notably, *Kat7* knockdown reduced cytosolic mtDNA levels (**Fig. 4C**). Consistently, it led to reduced phosphorylation of TBK1 (Ser172, p-TBK1) and IRF3 (Ser396, p-IRF3) (**Fig. 4D-G**), as well as diminished IL-1β production (**Fig. 4H**), indicating attenuated cGAS-STING activation and NLRP3 inflammasome signaling. Importantly, CMPK2 overexpression restored cytosolic mtDNA levels and innate immune signaling in *Kat7* deficient cells, whereas the catalytically inactive mutant failed to do so (**Fig. 4C-H**). Together, these results demonstrate that KAT7 orchestrates CMPK2-dependent mtDNA synthesis and release, establishing it as a critical epigenetic regulator of innate immune pathways during microglial activation.

**Fig. 4.**
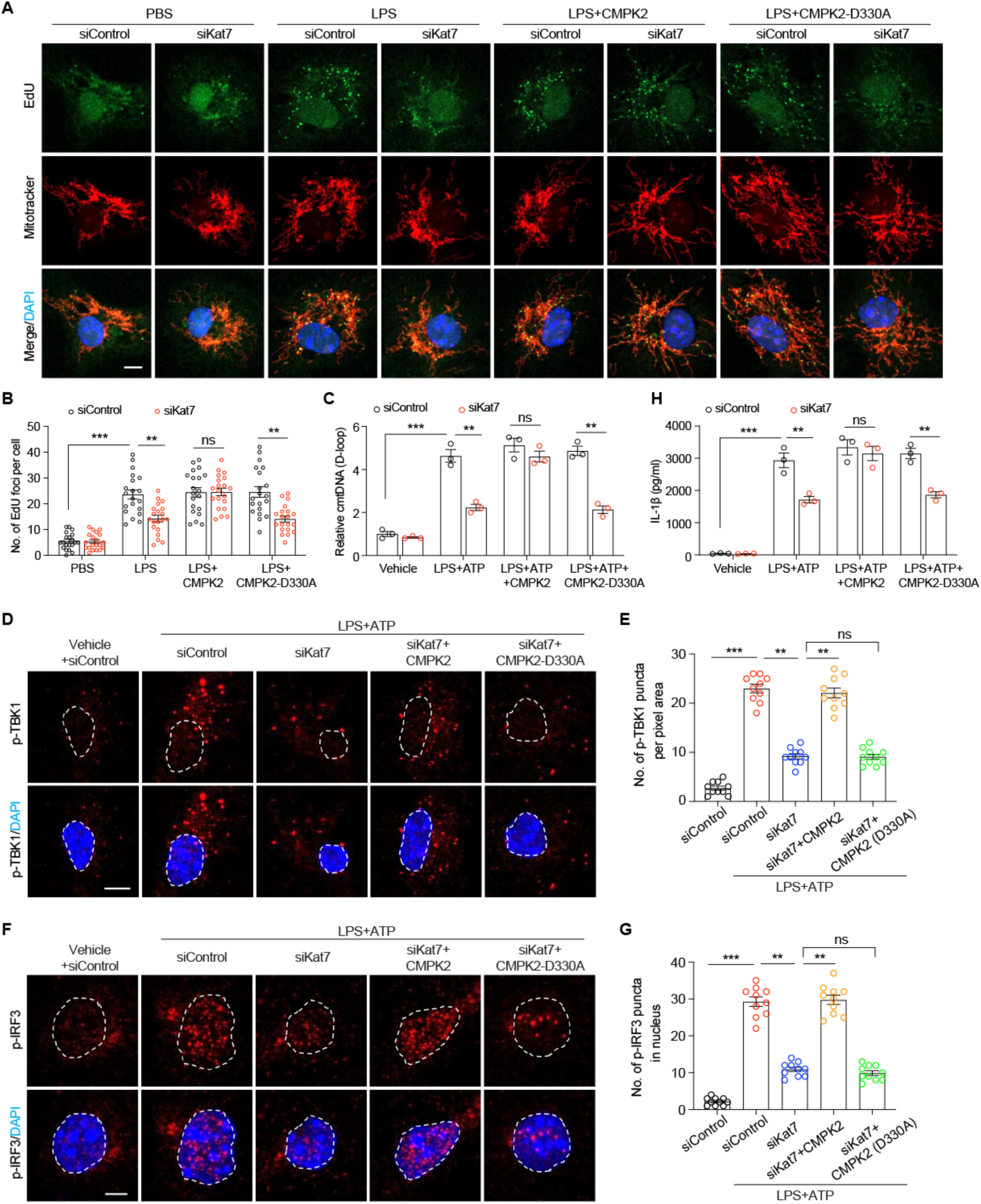
*Kat7* knockdown in microglia limits mtDNA replication and release through repressing *Cmpk2* transcription. **A-B**, Representative images (**A**) and quantification (**B**) of EdU-labelled newly synthesized mtDNA in primary microglia treated with LPS. Scale bar, 5 µm. n=20 cells from 3 replicates per group. **C**, Quantification of cytosolic mtDNA (cmtDNA) by qPCR (normalized to nuclear B2m DNA) in primary microglia treated with LPS plus ATP. D-loop indicated a specific fragment within mtDNA. n=3. **D-E**, Representative images (**D**) and quantification (**E**) of p-TBK1 in primary microglia treated with LPS plus ATP. Scale bar, 5 µm. n=10 cells from 2 replicates per group. **F-G**, Representative images (**F**) and quantification (**G**) of p-IRF3 in nuclei of primary microglia treated with LPS plus ATP. Scale bar, 3 µm. n=10 cells from 2 replicates per group. **H**, ELISA analysis of IL-1β secretion from primary microglia treated with LPS plus ATP. n=3. **p<0.01, ***p<0.001. One-way ANOVA test (**E** and **G**). Two-way ANOVA test (**B**, **C** and **H**). Data are presented as mean ± SEM.

### Microglia-specific *Kat7* deletion attenuates neuroinflammation, reduces Aβ pathology, and improves cognition in 5×FAD mice

Given our findings of elevated microglial KAT7 complex expression in AD, and its critical role in regulating neuroinflammation *in vitro*, we hypothesized that KAT7 promotes AD pathogenesis *in vivo* by driving cytosolic mtDNA-induced innate immune signaling. To test this, we generated *Kat7* floxed mice using the efficient additions with single-strand DNA inserts CRISPR (Easi-CRISPR) method^33^. Exon 2 of *Kat7* was flanked by loxP sites, and its deletion induces a frameshift mutation resulting in functional KO (**Suppl. Fig. 7**). These mice were crossed with Cx3cr1-CreER mice^34^ to achieve microglia-specific knockout (*Kat7* cKO) and subsequently bred with 5×FAD mice (**Fig. 5A**). Tamoxifen was administered at 2 months of age to avoid perturbing microglial development, and biochemical, pathological, and behavioral assessments were performed at 6 months. Microglia isolated from *Kat7* cKO mice exhibited efficient *Kat7* deletion (**Fig. 5B**). Notably, *Cmpk2*, which was markedly upregulated in 5×FAD microglia, was reduced by *Kat7* deletion (**Fig. 5C**). Given KAT7’s role in mtDNA synthesis and release, we assessed cytosolic mtDNA levels by qPCR analysis and found them elevated in 5×FAD microglia but significantly attenuated in *Kat7* cKO; 5×FAD mice (**Fig. 5D**). Correspondingly, phosphorylation of TBK1 and IRF3, key mediators of the cGAS-STING pathway, was markedly reduced in *Kat7* deficient microglia, along with decreased levels of the proinflammatory cytokines IL-1β and IL-6 (**Fig. 5E-H**). Further analysis revealed that microglial activation was suppressed in *Kat7* cKO; 5×FAD mice, as evidenced by a reduced number of Iba1-positive microglia in the hippocampus (**Fig. 5I**) and diminished protein levels of Iba1 and phosphorylated p65 (p-p65) (**Fig. 5J**), a key NF-κB effector downstream of cGAS-STING signaling^35^. These results demonstrate that microglial *Kat7* deletion suppresses neuroinflammation in 5×FAD mice.

**Fig. 5.**
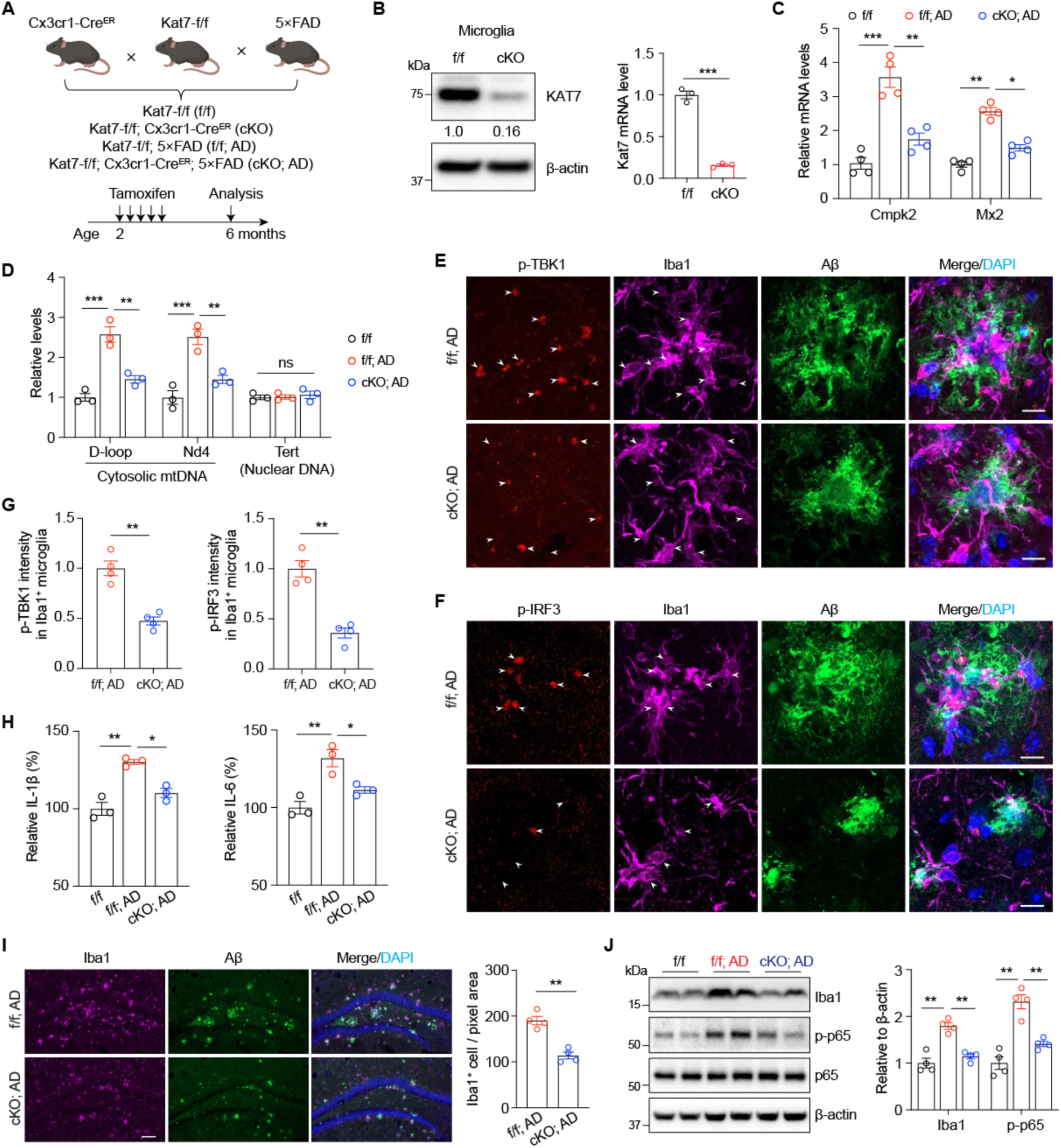
Microglia-specific *Kat7* deletion inhibits neuroinflammation in 5×FAD mice. **A**, Schematic diagram showing the strategy for generating *Kat7*-specific deletion in microglia in 5×FAD mice. **B**, Knockout efficiency of *Kat7* in microglia was examined by western blot (left) and qPCR analysis (right). Unpaired student’s t test. **C**, qPCR analysis of *Cmpk2* and *Mx2* in microglia isolated from 6-month-old mice. n=4. Two-way ANOVA test. **D**, Quantification of cytosolic mtDNA by qPCR (normalized to nuclear B2m DNA) in microglia isolated from 6-month-old mice. D-loop and Nd4 indicated a specific fragment within mtDNA. Tert and B2m indicated nuclear DNA. n=3. Two-way ANOVA test. **E-F**, Representative images of p-TBK1 (**E**) and p-IRF3 (**F**) staining in cortex region of 6-month-old AD mice. n=4 mice per group. Scale bar, 10 µm. One-way ANOVA test. **G**, Quantification of p-TBK1 (**E**) and p-IRF3 (**F**) intensity in microglia of 6-month-old AD mice. n=4 mice per group. Unpaired student’s t test. **H**, ELISA analysis of IL-1β and IL-6 production in cortex region of 6-month-old mice. **I**, Representative images (left) and quantification (right) of Iba1^+^ microglia in 6-month-old mice. n=4 mice per group. Scale bar, 100 µm. Unpaired student’s t test. **J**, Immunoblot analysis of Iba1 and p-p65 in the cortex of 6-month-old mice (left), and protein levels were normalized to β-actin (right). n=4 mice per group. Two-way ANOVA test. *p<0.05, **p<0.01, ***p<0.001. Data are presented as mean ± SEM.

Growing evidence indicates that microglia-driven neuroinflammation promotes Aβ deposition and disrupts synaptic activity, ultimately contributing to cognitive decline^5,36,37^. To determine whether *Kat7* deletion mitigates Aβ pathology, we performed Thioflavin S (TS) staining on brain sections from 6-month-old 5×FAD mice. Microglia-specific *Kat7* cKO mice exhibited a pronounced reduction in Aβ plaque burden across multiple brain regions, including the cortex and hippocampus (**Fig. 6A-B**). To evaluate synaptic plasticity, we performed field potential recordings to measure long-term potentiation (LTP) at Schaffer collateral-CA1 synapses in acute hippocampal slices. The LTP deficits characteristic of 5×FAD mice were significantly rescued by microglial *Kat7* deletion (**Fig. 6C-D**). We next assessed hippocampus-dependent spatial learning and memory using the Morris water maze. In line with restored synaptic function, *Kat7* cKO; 5×FAD mice demonstrated accelerated learning during training (**Fig. 6E**) and superior memory retention in the probe trial, spending more time in the target quadrant and crossing the former platform location more frequently compared to 5×FAD controls (**Fig. 6F-H**). Notably, swimming speed was comparable among groups (**Fig. 6I**), ruling out motor deficits as a confounding factor. Collectively, these results demonstrate that microglial *Kat7* deletion reduces Aβ accumulation, restores synaptic function, and improves cognitive performance in 5×FAD mice.

**Fig. 6.**
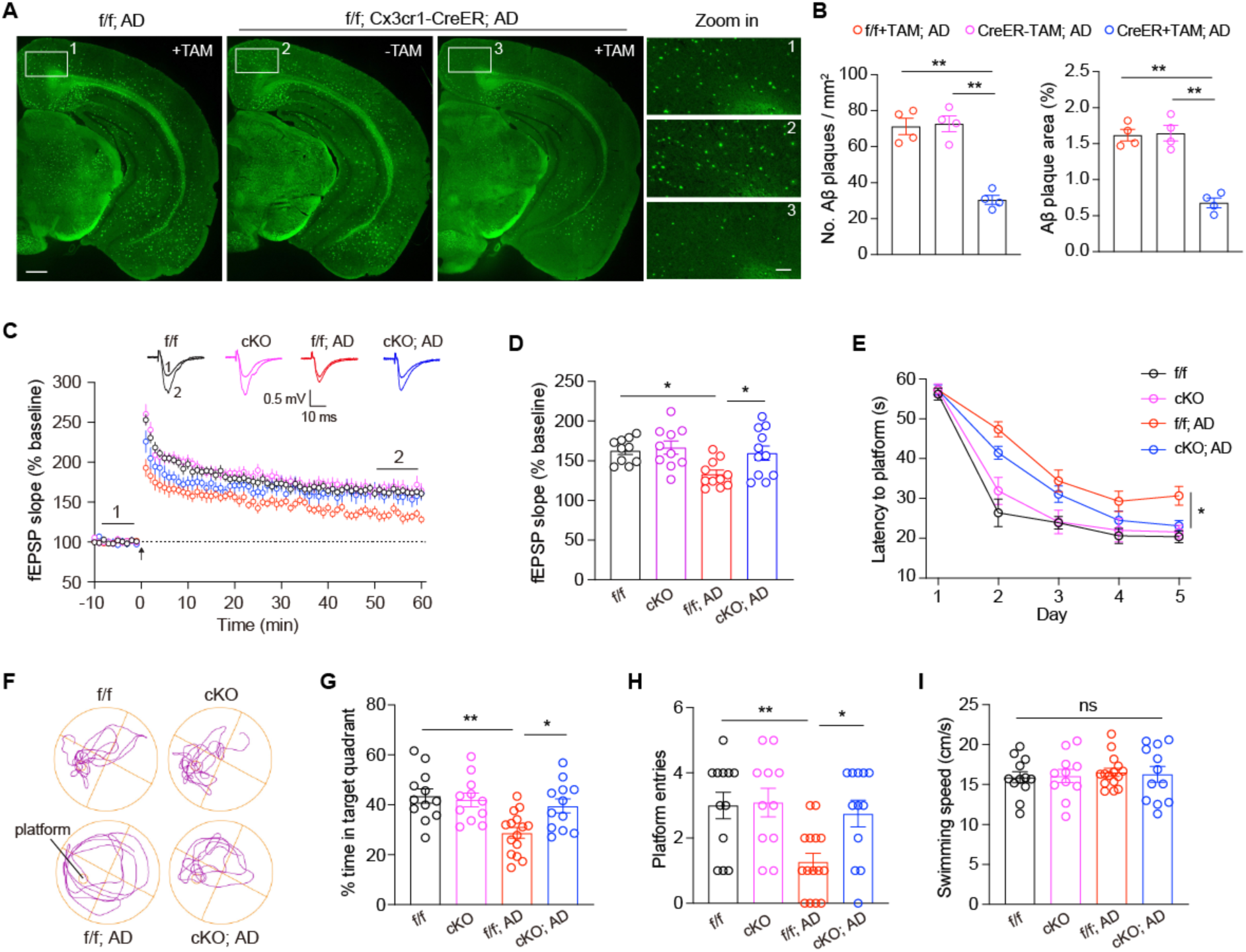
Microglia-specific *Kat7* deletion ameliorates Aβ pathology and improves cognitive function in 5×FAD mice. **A**, Representative images of TS staining in the brain sections of 6-month-old AD mice. Scale bar, 0.5 mm (left), 0.1 mm (right). TAM: tamoxifen. **B**, Quantification of TS-positive Aβ plaque number and area in 6-month-old AD mice. n=4 mice per group. One-way ANOVA test. **C**, TBS-induced LTP at Schaffer collateral to CA1 synapses in 6-month-old mice. Arrow indicates LTP induction. Sample traces represent fEPSP taken before (1) and 50 min after (2) TBS. **D**, Averaged fEPSP slopes during 50 to 60 min after the stimulation. n=10-11 slices from 5 mice per group. One-way ANOVA test. **E**, Time spent before reaching the hidden platform during training days in the Morris water maze test. Two-way ANOVA test. **F-I**, Representative traces (**F**), time spent in target quadrant (**G**), entries into platform zone (**H**) and swimming speed (**I**) during the probe test. n=11-15 mice per group. One-way ANOVA test. *p<0.05, **p<0.01. ns, nonsignificant. Data are presented as mean ± SEM.

### Pharmacological inhibition of KAT7 mitigates neuroinflammation, reduces Aβ burden, and improves cognition in 5×FAD mice

To determine whether pharmacological inhibition of KAT7 confers protection against AD, we first examined the effects of the KAT7 inhibitor WM-3835 in primary mouse microglia treated with oligomeric Aβ42. Consistent with our knockdown results (**Fig. 3J** and **Fig. 4A-B**), WM-3835 treatment suppressed Aβ-induced *Cmpk2* upregulation and markedly reduced CMPK2-dependent mtDNA synthesis (**Fig. 7A-B**). Before evaluating KAT7 inhibition *in vivo*, we examined whether *Kat7* deletion impacts brain function beyond early development, given its neuronal expression and essential roles in neural stem cell differentiation and cortical development^29^. We generated excitatory neuron-specific *Kat7* cKO mice using CamKII-Cre (**Suppl. Fig. 8A-B**), which induces deletion beginning at 2-3 weeks of age^38^. Unlike mice with embryonic *Kat7* deletion^29^, these cKO mice were viable, healthy, and fertile. Electrophysiological recordings from hippocampal CA1 pyramidal neurons in acute slices revealed normal miniature excitatory postsynaptic current (mEPSC) amplitude and frequency, as well as intact LTP at Schaffer collateral-CA1 synapses (**Suppl. Fig. 8C-D**), indicating preserved synaptic transmission and plasticity. These results suggest that loss of KAT7 is well tolerated in mature neurons. We then evaluated the therapeutic potential of WM-3835 *in vivo*. Five-month-old 5×FAD mice received intracerebroventricular (ICV) infusion of WM-3835 for four weeks via an osmotic pump (**Fig. 7C**). Post-treatment analysis confirmed robust target engagement, evidenced by a pronounced reduction in H3K14ac levels across the brain (**Fig. 7D-E**). Notably, WM-3835 significantly decreased microgliosis (**Fig. 7F**), lowered cytosolic mtDNA levels (**Fig. 7G**), and attenuated cGAS-STING pathway activation in microglia (**Fig. 7H**). Consistent with these effects, WM-3835 treatment also markedly reduced Aβ plaque burden in the hippocampus and cortex (**Fig. 7I**). Behaviorally, WM-3835-treated 5×FAD mice exhibited improved spatial learning and memory in the Morris water maze, with faster acquisition during training and superior performance in the probe trial (**Fig. 7J-L**). These cognitive benefits closely mirrored those observed in microglia-specific *Kat7* cKO mice. Together, our results demonstrate that pharmacological inhibition of KAT7 mitigates microglial activation, reduces Aβ pathology, and improves cognitive function in 5×FAD mice, providing preclinical evidence for WM-3835 as a potential therapeutic strategy for AD.

**Fig. 7.**
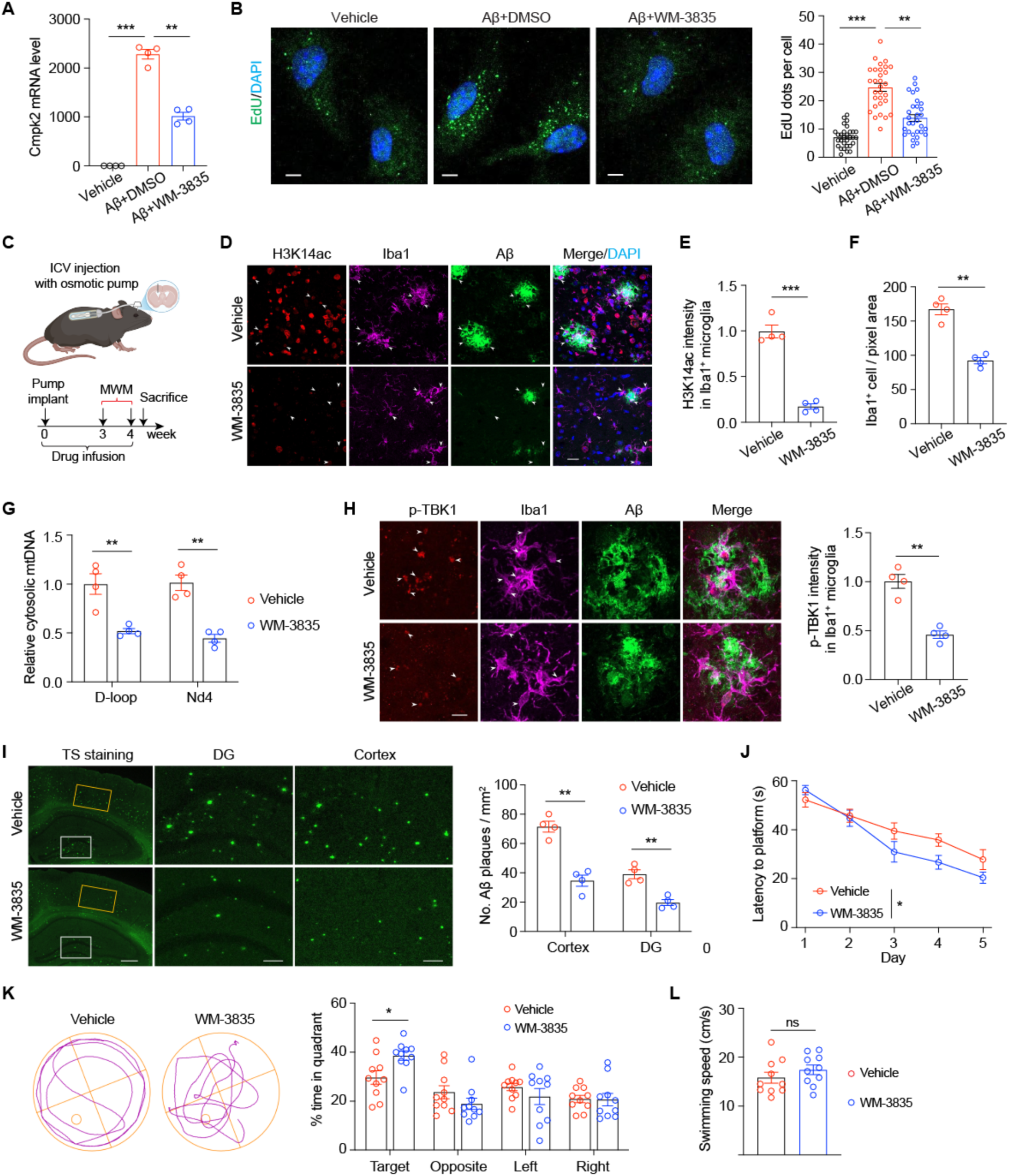
Pharmacological inhibition of KAT7 reduces neuroinflammation and Aβ burden in 5×FAD mice. **A**, qPCR analysis of *Cmpk2* levels in primary microglia treated with WM-3835 and oligomeric Aβ42. n=4. **B**, Representative images (left) and quantification (right) of EdU-labelled newly synthesized mtDNA in primary microglia treated with WM-3835 and Aβ42. Scale bar, 4 µm. n=30 cells from 3 replicates per group. **C**, Schematic diagram showing the strategy for delivering vehicle or WM-3835 into ICV via osmotic pump in 5-month-old 5×FAD mice. **D-F**, Representative images (**D**) and quantification of H3K14ac intensity in microglia (**E**) and Iba1^+^ microglia number (**F**) in cortex of 6-month-old AD mice treated with vehicle or WM-3835. Scale bar, 20 µm. n=4 mice per group. **G**, Quantification of cytosolic mtDNA by qPCR (normalized to nuclear B2m DNA) in microglia isolated from 6-month-old AD mice treated with vehicle or WM-3835. D-loop and Nd4 indicated a specific fragment within mtDNA. B2m indicated nuclear DNA. **H**, Representative images (left) and quantification (right) of p-TBK1 in microglia of 6-month-old AD mice treated with vehicle or WM-3835. Scale bar, 10 µm. n=4 mice per group. **I**, Representative images (left) and quantification (right) of TS staining in the cortex and hippocampus of 6-month-old AD mice treated with vehicle or WM-3835. Scale bar, 0.4 mm (left), 0.1 mm (middle and right). n=4 mice per group. **J**, Time spent before reaching the hidden platform during training days in the Morris water maze test. **K-L**, Time spent in different quadrants (**K**) and swimming speed (**L**) during the probe test. n=10 mice per group. *p<0.05, **p<0.01, ***p<0.001. Unpaired student’s t test (**E**, **F**, **H** and **L**). One-way ANOVA test (**A-B**). Two-way ANOVA test (**G**, **I**, **J** and **K**). Data are mean±SEM.

## DISCUSSION

Cytosolic mtDNA has emerged as a potent driver of neuroinflammation through activation of innate immune signaling. Here, we identify the histone acetyltransferase KAT7 as a critical upstream regulator of this mtDNA-initiated inflammatory cascade in microglia (**Fig. 8**). Mechanistically, KAT7 promotes H3K14 acetylation at the *Cmpk2* promoter during microglial activation, thereby enhancing *Cmpk2* expression, increasing mtDNA synthesis and release, and fueling downstream inflammatory responses. Multiple lines of evidence indicate that this pathway is active in AD. First, the expression of KAT7 complex and its histone mark H3K14ac are elevated in microglia from 5×FAD mice and human AD brains. Second, Aβ oligomers induce *Cmpk2* expression and mtDNA replication in primary microglia, processes that depend in part on KAT7 expression or activity. Third, *Cmpk2* expression and cytosolic mtDNA levels are elevated in microglia from 5×FAD mice, both of which are reduced upon *Kat7* deletion. Finally, both microglia-specific *Kat7* deletion and pharmacological inhibition suppress innate immune signaling and neuroinflammation, leading to reduced Aβ burden and improved cognitive function in 5×FAD mice. Our findings thus reveal an epigenetic-mitochondrial axis that mechanistically links Aβ deposition to chronic immune activation. Disrupting this self-perpetuating loop offers a potential therapeutic strategy for AD and related neurodegenerative disorders.

**Fig. 8.**
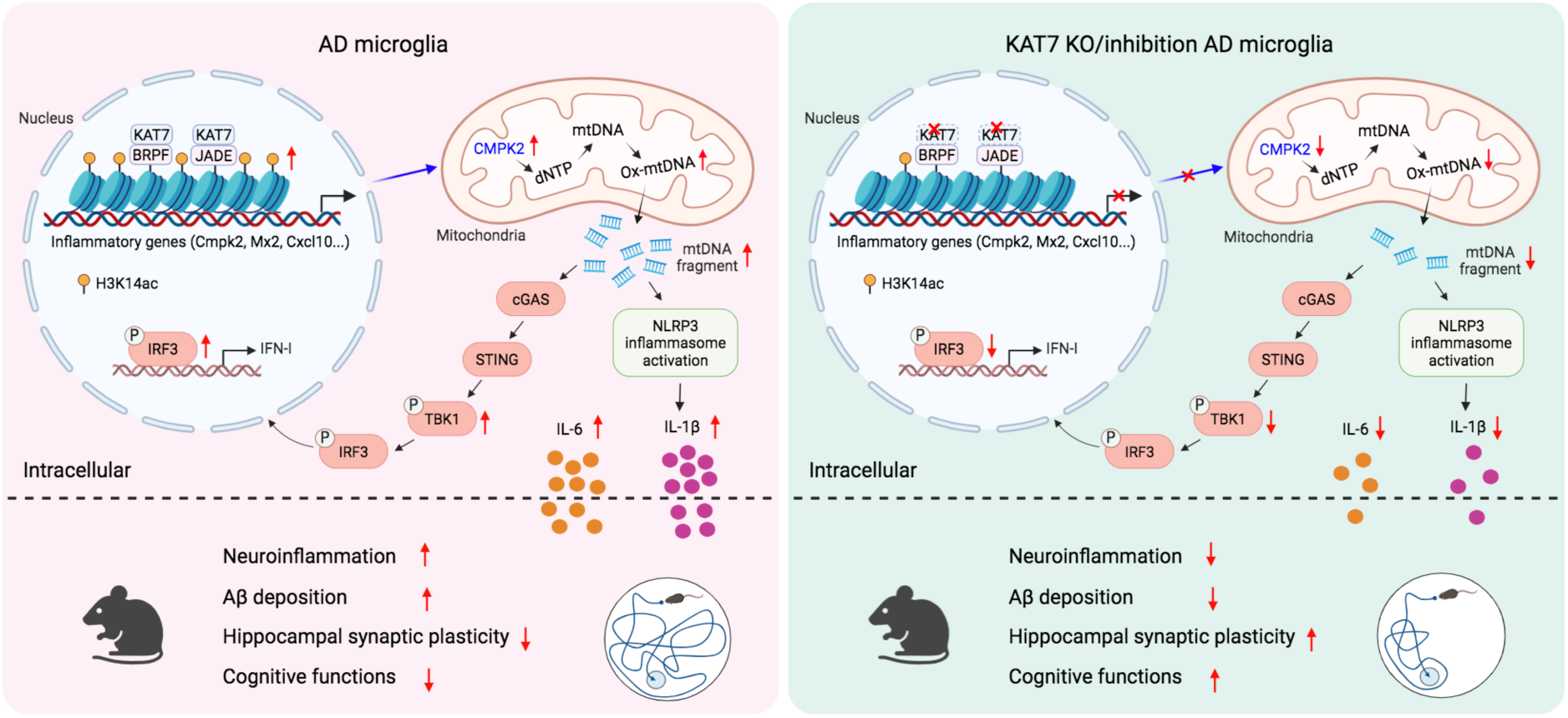
Summary diagram. Our results support a model in which KAT7 acts as a central epigenetic driver of neuroinflammation by promoting H3K14ac-dependent *Cmpk2* expression. Through this mechanism, KAT7 integrates nuclear and mitochondrial responses to amplify neuroinflammation, thereby contributing to Aβ accumulation and synaptic dysfunction in AD.

Microglial activation requires extensive metabolic reprogramming to meet heightened bioenergetic and biosynthetic demands^39,40^. KAT7-meidated upregulation of CMPK2, a key enzyme for mtDNA replication, may help fulfill these needs. However, while essential for sustaining microglial activation, excessive mtDNA replication renders mitochondria vulnerable to stress-induced mtDNA release, thereby amplifying innate immune signaling. Notably, a similar CMPK2-mtDNA pathway drives cGAS-STING and NLRP3 activation in macrophages during inflammation^30,32,41^, underscoring its conserved role in innate immunity across myeloid lineages. Future studies will be required to directly define the role of CMPK2 and its downstream mtDNA replication in neuroinflammation and AD. Of note, a recent study showed that CMPK2 is also upregulated in microglia during ischemic stroke, where it promotes neuroinflammation and brain injury^42^. Therefore, the KAT7-CMPK2 pathway may represent a general mechanism by which microglia couple metabolic reprogramming to innate immune activation across diverse chronic inflammatory conditions.

Histone acetylation in neurons has long been associated with learning and memory^43-45^, and several altered histone acetylation marks have been reported in AD mouse models and patients^46-48^. However, the functional relevance of individual histone modifications in AD remains poorly defined. Here, we show that KAT7-dependent H3K14ac is low in microglia from WT mice and healthy controls but markedly and specifically elevated in AD brains. Functional studies further reveal a microglia-specific pathogenic role for KAT7 in 5×FAD mice. These findings highlight the importance of examining histone acetylation and other epigenetic modifications in a cell type-specific manner. Notably, KAT7 has also been implicated in ageing and cellular senescence, where it enhances transcription of the cyclin-dependent kinase inhibitor *p15^INK4b^* to promote cell cycle arrest^49^. Downregulation of *Kat7* in aged mice reduces hepatocyte senescence, alleviates liver inflammation, and extends lifespan^49^. Given its beneficial effects in both AD and aging contexts, KAT7 inhibition may represent a promising strategy to mitigate neurodegeneration and promote brain resilience across the lifespan.

Our data establish CMPK2 as a key target of KAT7, but additional downstream genes, including several interferon-responsive genes, are also likely to contribute to chronic neuroinflammation. While KAT7 is best known for catalyzing H3K14ac, it also mediates other histone acylations, such as propionylation and crotonylation^50^, and can modify both alternative histone sites and non-histone substrates^23,51^. These activities may further diversify its regulatory functions and shape the transcriptional landscape of activated microglia. Regardless, pharmacological inhibition of KAT7 is expected to suppress all of these activities, providing a unified strategy to blunt its proinflammatory effects in AD. While the 5×FAD model used in this study recapitulates robust Aβ pathology, it does not develop tau pathology, a defining feature of human AD. Future studies employing tauopathy or mixed pathology models will therefore be essential to fully delineate the role of KAT7 in AD pathogenesis. Nevertheless, our findings identify the KAT7-CMPK2 axis as a critical regulator of microglial mitochondrial immunity, linking epigenetic control to neuroinflammation and AD progression. These insights highlight the epigenetic-mitochondrial axis as a promising therapeutic target not only for AD but also for other neurological diseases characterized by chronic neuroinflammation.

## MATERIALS AND METHODS

### Human brain samples

Formalin-fixed paraffin-embedded (FFPE) human postmortem brain samples from AD patients and controls (aged-matched and died of certain cause unrelated to dementia) were obtained from Johns Hopkins Brain Resource Center. Subject demographics were listed in **Tables S1**. The study using patient samples/data was approved by the Johns Hopkins University School of Medicine Office of Human Subjects Research Institutional Review Boards.

### Mice

All procedures related to animal care and treatment were approved by the Johns Hopkins University Animal Care and Use Committee and met the guidelines of the National Institute of Health Guide for the Care and Use of Laboratory Animals. All animals were group housed in a standard 12-hour light/dark cycle with ad libitum access to food and water. The following mouse lines were used for the experiments: C57BL/6J (Jackson Laboratory, 000664), 5×FAD (Jackson Laboratory, 008730), Cx3cr1-Cre^ER^ (Jackson Laboratory, 021160), Camk2a-Cre (Jackson Laboratory, 005359). *Kat7* floxed mice were generated at Transgenic Core of Johns Hopkins University. Male mice were used for all experiments unless otherwise noted. Female mice were also used for the behavioral tests. Mice with *Kat7* specific knockout in microglia were induced by tamoxifen (S1238, Selleck). Briefly, tamoxifen was dissolved in corn oil (C8267, Sigma) to a final concentration of 20 mg/ml. 2-month-old mice were administered tamoxifen via intraperitoneal injection at 100 mg/kg for five consecutive days.

### Generation of *Kat7* floxed mice

*Kat7* floxed mice were generated at Transgenic Core of Johns Hopkins University using the Easi-CRISPR method, as previously described^33^. Two single-guide RNAs (sgRNAs) were designed by http://crispor.tefor.net/. The sequences were as follows: sgRNA #1 (reverse strand), AAGTACCAAGTTCCAACATAAGG; and sgRNA #2 (forward strand), GATACTGCTCCTGAGCTTGATGG. Two crRNAs containing each sgRNA and ssDNA donor containing the homology arms and floxed exon sequences were custom synthesized from IDT company. The annealed crRNA and tracrRNA (IDT) were diluted in microinjection buffer (0.25 mM EDTA and 10 mM Tris-HCl, pH 7.4) and mixed with Cas9 protein (30 ng/µl; IDT) to obtain ctRNP complexes. One-cell embryos of C57BL/6J mice were microinjected with a mixture of floxing ssDNA donors and two ctRNP complexes and were transferred into the oviducts of pseudopregnant ICR females (Charles River Laboratories). Successful insertions of two LoxP sites were detected by PCR genotyping of mouse tails and confirmed by Sanger sequencing. The primers used are provided in **Table S2**.

### Primary microglial cell preparation and stimulation

Primary microglial cells were prepared from neonatal mice (day 0-2). Briefly, brain tissues were quickly removed, and the meninges were carefully stripped in ice-cold HBSS. The cortex and hippocampi were then digested with 0.25% trypsin (Quality Biological) at 37 ℃ for 15 min and gently pipetted to generate single cells with DMEM containing 10% heat-inactivated fetal bovine serum (FBS; Avantor) and 1% penicillin/streptomycin (P/S; Quality Biological), followed by plating on poly-D-lysine-coated T75 flasks. After 11-14 days, primary microglia were separated from the mixed glial culture using a shake-off method (90 rpm for 2 hours). The collected microglia were seeded in the poly-D-lysine-coated plates and its purity was confirmed by Iba1 immunostaining.

For LPS stimulation, microglial cells were cultured in basic DMEM without FBS and treated with LPS (0.2 µg/ml, Sigma) for 6 h. For the assessment of mtDNA release, microglia were primed with LPS (0.2 µg/ml, Sigma) for 6 h, followed by ATP (2 mM, Sigma) treatment for 30 min. For oligomeric Aβ42 stimulation, microglial cells were cultured in basic DMEM without FBS and treated with oligomeric Aβ42 (1 µM, rPeptide) for 6 h or 24 h.

### Isolation of microglia from adult mouse brain

6-month-old mice were anesthetized with isoflurane and perfused transcardially with cold saline. Brain tissue was freshly harvested, cut into small pieces, and digested with collagenase (Type IV, 5 mg/ml, Sigma) and DNase I (50 µg/ml, Sigma) for 1 hour at 37 ℃ with 250 rpm. The digested brain tissues were transferred to a 15 ml Dounce homogenizer and homogenized gently on ice. Brain tissue homogenates were suspended in HBSS, filtered with cell strainers (70 µm), and centrifuged at 500g for 5 min (4 °C) to collect the cell pellets. Then, 90% Percoll solution was prepared using absolute Percoll (Cytiva) and 10× HBSS (9:1, v/v), and further diluted (v/v) to 70, 37, and 30% with 1× HBSS. Cell pellets were suspended in a 37% Percoll solution. Microglia were isolated by density gradient centrifugation. Density gradient was added into 15 ml tubes, by layers of Percoll solution from bottom to top containing: 70%, 37%, and 30% Percoll solution and HBSS. Centrifugation was carried out in a horizontal centrifuge at 2000g for 30 min (4 °C). Microglia were converged on the interphase between the 37% and 70% Percoll solution. Isolated microglia were washed with 10× volumes of PBS and centrifuged at 500g for 5 min (4 °C). Microglia was further purified by CD11b MicroBeads (Miltenyi Biotec, 130-093-634) according to the manufacturer’s protocol.

### BV2 cell culture

The mouse microglial BV2 cell line was a gift from Dr. Tony Wyss-Coray’s laboratory at Stanford University^52^. Cells were cultured in DMEM supplemented with 10% FBS and 1% penicillin/streptomycin and maintained in an incubator at 37 °C with 5% CO_2_. Adherent cells were split using 1× TrypLE (Gibco).

KAT7-KO BV2 cells were generated using CRISPR-Cas9 method. Guide RNA (GACTCGGGCAGATCGGCGCG) targeting mouse Kat7 was cloned into LentiCRISPR-v2-Puro (Addgene, #98290). The primers used to design the single-guide RNA (sgRNA) targets were (5’ to 3’) Kat7 forward CACCGGACTCGGGCAGATCGGCGCG and Kat7 reverse AAACCGCGCCGATCTGCCCGAGTCC. Lentiviral particles containing Kat7 sgRNA were packaged using the 3rd generation lentivirus system and used to infect BV2 cells. One day after infection, the medium was changed to fresh DMEM containing 10% FBS and 1% P/S. Cells were then treated with puromycin (4 µg/ml) for 5 days to select for successfully transduced cells. Single clones were obtained using limiting dilution and were analyzed by western blotting and Sanger sequencing to confirm KAT7 deletion. A scrambled gRNA control was also used as negative control (5’-GCGCCAAACGTGCCCTGACG-3’).

For the overexpression in BV2 cells, the GFP in the lentivirus vector pLenti-EF1a-GFP-P2A-Puro (a gift from Dr. Shuying Sun lab at Johns Hopkins University) was replaced by mouse CMPK2, CMPK2-D330A, human KAT7, KAT7-E508Q and JADE2 DNA fragments at the AgeI and BamHI site using NEBuilder HiFi DNA Assembly Cloning Kit (New England Biolabs, #E5520). Lentiviral particles were packaged using the 3rd generation lentivirus system and used to infect BV2 microglia. Cells were then treated with puromycin (4 µg/ml) to select the transduced cells.

### siRNA transfection

siRNAs were purchased from Dharmacon and transfected into primary microglia with the final concentration of 40 nM using Lipofectamine RNAiMAX (Invitrogen) according to the manufacturer’s instructions. The sequences of siRNA used in this study are as follows: siControl sense, UGGUUUACAUGUCGACUAA; mouse Kat7 siRNA#1 sense, GAACCGAAGAUUCCGAUUU; siRNA#2 sense, UGUUUGAAGUAGACGGCAA.

### Enzyme-linked immunosorbent assay (ELISA)

The collected cultured medium of BV2 cells or primary microglia was centrifuged at 500 g for 5 min (4°C) and the supernatant was processed for analysis with mouse IL-6 (431301, BioLegend) and IL-1β (432601, BioLegend) ELISA kits, according to the manufacturer’s instructions. For cortical IL-6 and IL-1β detection, the cortex tissues were removed from indicated mice and rapidly immersed in RIPA buffer (Sigma) containing protease inhibitor cocktails (Roche). Total IL-6 and IL-1β protein levels were measured by the ELISA kit and normalized first to total protein level quantified by the Pierce BCA Protein Assay Kit (Thermo) and then to the floxed (WT) group.

### Western blotting

Proteins were isolated from cultured cells or brain tissues with RIPA buffer (Sigma) containing protease inhibitor cocktails (Roche). Samples were separated on Novex Tris-Glycine Mini Protein Gels (4 to 20%, Invitrogen) and transferred to nitrocellulose membranes (Bio-Rad), which were incubated with appropriate antibodies for overnight at 4°C. Primary antibody concentrations were as follows: anti-KAT7 (rabbit, 1:1000, Cell Signaling Technology, #58418), anti-JADE2 (rabbit, 1:2000, Proteintech, 11513-1-AP), anti-iNOS (rabbit, 1:3000, GeneTex, GTX130246), anti-Iba1 (rabbit, 1:1000, Wako, #019-19741), anti-p65 (rabbit, 1:1000, Cell Signaling Technology, #8242), anti-phospho-p65 (Ser536) (rabbit, 1:1000, Invitrogen, #MA5-15160), anti-CMPK2 (rabbit, 1:1000, Novus Biologicals, # NBP1-80653), anti-GAPDH (mouse, 1:5000, Proteintech, 60004-1-lg), anti-β-actin (mouse, 1:5000, Proteintech, 66009-1-lg), anti-H3K14ac (rabbit, 1:2000, Millipore, #07-353), anti-H3 (rabbit, 1:3000, Proteintech, 17168-1-AP). After wash, the membranes were incubated horseradish peroxidase (HRP)-conjugated secondary antibody (Cytiva, 1:5000). Immunoreactive bands were visualized using Western Chemiluminescent HRP Substrate (Millipore, #WBKLS0500) and analyzed with ImageJ.

### Real-time qPCR

Total RNA was isolated from samples with TRIzol reagents (Invitrogen) and was reverse transcribed into cDNA using the HiScript III RT SuperMix for qPCR (+gDNA wiper) kits (R323-01, Vazyme). Relative quantitation was determined using the QuantStudio 6 Flex detection system (Applied Biosystems) that measures real-time SYBR green fluorescence and then calculated by means of the comparative Ct method (2^−ΔΔCt^) with the expression of Gapdh or β-actin as an internal control. The sequences of primers used are provided in **Table S3**.

### EdU staining

To measure newly synthesized mtDNA in primary microglia, in the presence of 10 µM EdU, the cells were treated with LPS (0.2 µg/mL) or oligomeric Aβ42 (1 µM) for 6 hours and then were incubated with MitoTracker (250 nM, Invitrogen) for 30 minutes. After cell fixation, permeabilization and blocking, EdU staining was performed according to the manufacturer’s protocol using a Click-iT EdU Alexa Fluor 488 Imaging Kit (Invitrogen). The nucleus was stained with DAPI for 5 minutes. Images were collected with a Zeiss LSM 900 confocal microscope and analyzed using ImageJ software (NIH).

### Immunofluorescence

For cells immunofluorescence, primary microglia were washed with PBS and fixed with 4% PFA for 20 minutes at room temperature. After PBS wash, cells were permeabilized with 0.2% Triton X-100 and blocked with blocking buffer (2% donkey serum plus 1% BSA in PBS). Cells were incubated with primary antibodies overnight at 4°C. On the next day, cells were washed with PBS and incubated with secondary antibodies (1:100, Jackson ImmunoResearch) for 1 hour at room temperature.

For immunofluorescence of mouse brain cryosections, anesthetized mice were perfused transcardially with PBS, followed by 4% cold PFA in PBS. Brains were removed and fixed in 4% PFA at 4°C overnight. After dehydration by 30% sucrose, brains were embedded in OCT (Tissue-Tek) and cut into 30-µm-thick sections on cryostat microtome (Leica). Sections were permeabilized and blocked with 0.3% Triton X-100 and 5% donkey serum in PBS for 1 hour at room temperature, and incubated with primary antibodies at 4°C overnight. After washing three times with PBS, slices were incubated with secondary antibodies (1:100, Jackson ImmunoResearch) for 2 hours at room temperature.

For immunofluorescence of formalin-fixed paraffin-embedded (FFPE) human patient tissues, the brain tissue sections were deparaffinized in a 60°C oven for 2 hours, followed by xylene washes twice, each for 10 minutes at room temperature. Tissues were then rehydrated in a graded series of ethanol washes. Slides were rinsed with deionized water twice and transferred into sodium citrate buffer (10 mM, adjust pH to 6.0) for antigen retrieval at 120 °C for 20 minutes. After cooling down to room temperature, sections were washed with PBS and permeabilized with 0.3% Triton X-100 in PBS for 20 minutes at room temperature, then blocked with blocking buffer (20% donkey serum plus 1%BSA in PBS) for 1 hour at room temperature, and incubated with primary antibodies at 4°C overnight. On the next day, samples were incubated with secondary antibodies (1:100, Jackson ImmunoResearch) for 2 hours at room temperature. After washing three times with PBS, tissues were incubated with 0.1% Sudan Black B in 70% ethanol for 30 minutes to quench autofluorescence.

After samples were stained with DAPI and washed with PBS, samples were mounted using an aqueous mounting medium (Aqua-Poly/Mount, Polysciences). Images were obtained with Zeiss LSM900 confocal microscope and analyzed with ImageJ. Primary antibody concentrations were as follows: anti-Iba1 (goat, 1:200, Novus Biologicals, NB100-1028), anti-H3K14ac (rabbit, 1:100, Millipore, #07-353), anti-Aβ (mouse, 1:200, Biolegend, #803004), anti-phospho-TBK1 (rabbit, 1:100, Cell Signaling Technology, #5483), anti-phospho-IRF3 (rabbit, 1:100, Cell Signaling Technology, #4947), anti-GFAP (mouse, 1:400, Invitrogen, 14-9892-82), anti-NeuN (mouse, 1:200, Millipore, MAB377).

### Preparation of oligomeric Aβ_1-42_

HFIP (hexofluoro-isopropanol) treated human Aβ_1-42_ peptides (rPeptide, A-1163) were first dissolved in dimethyl sulfoxide (DMSO) to a final concentration of 5 mM, and this solution was then diluted with cold phenol-free basal culture media to a final concentration of 250 µM. Oligomeric Aβ_1-42_ was prepared by incubation for 24 h at 4 °C, and this solution was aliquoted and stored at -80 °C before use.

### Thioflavin S (TS) staining

Aβ plaques were labeled by Thioflavin S staining on brain sections that were stained with 0.01% thioflavin S (T1892, Sigma) in 50% ethanol for 10 min. Then, sections were washed twice with 50% ethanol and three times with PBS. Brain sections were mounted for imaging and analyzed using a Zeiss LSM 900 confocal microscope.

### Measurement of cytosolic mtDNA

Microglia were resuspended in digitonin buffer (50 mM HEPES, pH 7.4, 150 mM NaCl, and 25 µg/ml digitonin) and incubated for 10 min at room temperature, followed by centrifugation at 2000g for 10 min at 4 °C. The supernatant containing cytosolic mtDNA (cmtDNA) was used for qPCR. The pellet was used for nuclear DNA extraction with QIAamp DNA Mini Kit (Qiagen) according to the manufacturer’s instructions. The cmtDNA in the supernatant was normalized to the nuclear DNA (B2m gene) in the pellet for each sample. D-loop and Nd4 were used to assess mtDNA expression. B2m and Tert was used to assess nuclear DNA expression. The sequences of primers used are provided in **Table S4**.

### RNA sequencing

Three biological replicates were sequenced per group. For each sample, RNA was extracted from BV2 cells with TRIzol reagents (Invitrogen). High-throughput RNA sequencing (RNA-seq) was performed by Illumina NovaSeq 6000 at Novogene (CA, USA). The raw sequencing data were aligned to the mouse preference genome (GRCm39, mm39) using HISAT2 (v2.0.5). Reads on each GENCODE annotated gene were counted using HTSeq, and then differential gene expression analysis was performed using DESeq2 R package. GO pathway analysis was conducted with DAVID tools (https://davidbioinformatics.nih.gov/).

### Cleavage Under Targets & Tagmentation (CUT&Tag)

CUT&Tag was performed with Hyperactive Universal CUT&Tag Assay Kit for for Illumina Pro (TD904, Vazyme) according to the manufacturer’s instructions. In brief, BV2 cells were collected and counted same number for each group. Nuclei were isolated from BV2 cells and bounded to Concanavalin A (ConA)-coated beads. Subsequently, Nuclei were resuspended in antibody buffer and incubated with primary antibodies against H3K14ac (rabbit, Cell Signaling Technology, #7627) and IgG (rabbit, Cell Signaling Technology, #66362) at 4 °C overnight. On the next day, samples were incubated with goat anti-rabbit secondary antibodies (1:50, Cell Signaling Technology, #35401). The samples were incubated with pA/G-Tn5 transposase. After transposon activation and fragmentation, 0.5 pg Spike-in DNA was added to each sample and total DNA was isolated, amplified, and purified to construct library. The library for sequencing was constructed with TruePrep Index Kit V2 for Illumina (TD202, Vazyme) and VAHTS DNA Clean Beads (N411, Vazyme) were used for purification steps. The library was sequenced on an Illumina NovaSeq (PE 150) at Novogene. Raw sequencing reads were trimmed using Cutadapt 5.0 (https://cutadapt.readthedocs.io/en/stable/). Trimmed reads were then aligned to the mouse reference genome mm10 with the Spike-in sequence using Bowtie2 (version 2.3.5.1). Bam files with low-quality reads were filtered and duplicates were removed using Samtools v1.18. Reads were then normalized to Spike-in using Bedtools v2.31.0. Peaks were then called with SEACR v1.3. Differential peak analysis was analyzed by MAnorm2 and annotated by CHIPseeker with a p<0.05 cutoff.

### ChIP-qPCR

ChIP experiments were performed according to the procedure described previously^53^. BV2 cells were fixed with 1% formaldehyde for 15 min at room temperature. The fixed cells were lysed in lysis buffer (1% SDS, 5 mM EDTA, 50 mM Tris-HCl, pH 8.1) containing protease inhibitor cocktail. The lysates were then sonicated to generate chromatin fragments of ∼500 bp in length. Cell debris was removed by centrifugation and supernatant were collected. A dilution buffer (150 mM NaCl, 2 mM EDTA, 1% Triton X-100, and 20 mM Tris-HCl, pH 8.1) containing protease inhibitor cocktail was subsequently applied (1:9 ratio) and the chromatin solution (40 µl aliquot as the input) was then incubated with specific antibodies (2 µg) at 4°C overnight with mild rotation. 30 µl Protein A magnetic beads (Invitrogen) were added for incubation of 2 hours. Beads were sequentially washed with the following buffers: TSE I (150 mM NaCl, 2 mM EDTA, 0.1% SDS, 1% Triton X-100, 20 mM Tris-HCl, pH 8.1), TSE II (500 mM NaCl, 2 mM EDTA, 0.1% SDS, 1% Triton X-100, 20 mM Tris-HCl, pH 8.1), buffer III (0.25 M LiCl, 1% Nonidet P-40, 1 mM EDTA, 1% sodium deoxycholate, and 10 mM Tris-HCl, pH 8.1), and Tris-EDTA buffer. The input and the precipitated DNA-protein complex were de-crosslinked at 65°C for 12 hours in elution buffer (1% SDS, 0.1 M NaHCO_3_) with RNase A and Proteinase K. Then DNA was purified using QIAquick PCR Purification Kit (Qiagen). Quantification of the precipitated DNA fragments were performed with real-time PCR using primers listed in **Table S5**.

### Intracerebroventricular (ICV) injection

5-month-old 5×FAD mice were used for ICV injection of WM-3835 (S9805, Selleck). Briefly, 20 mice were randomly separated into two groups (WM-3835 and vehicle, 10 mice per group), then deeply anesthetized with isoflurane and immobilized using a stereotactic device. To implant osmotic pumps in the mice, osmotic pumps (1004W for 4 weeks infusion, RWD) matched with Brain infusion kit (Bic-3, RWD) were loaded according to the manufacturer’s instructions. 100 µL of vehicle (5% DMSO, 40% PEG300, and 55% saline) or WM-3835 (1 mM) was filled in the osmotic pump, and the Bic-3 kit/tubing (2 cm) was backfilled before the two parts were connected. In addition to the incision on the scalp, the pocket for the osmotic pump was obtained by stretching the space between the skin and the muscle in the back with sterile forceps. The detachable top part of the infusion cannula was held with a holder. A 0.5-1 mm burr hole was drilled in the skull, and the cannula tip was gently implanted into the lateral ventricle (coordinates, bregma: anterior/posterior, -0.5 mm; medial/lateral, 1.0 mm; and dorsal/ventral: -2.3 mm). The osmotic pump was slowly positioned in the pocket under the back skin simultaneously. The position of the cannula was secured with instant adhesives, and the skin was sutured with suture thread. ICV infusion was performed for four weeks. Then, the mice were sacrificed and processed for pathology analyses.

### RNAscope *in situ* hybridization

Fixed brains were embedded in OCT (Tissue-Tek) and sectioned at a thickness of 14 µm. RNAscope Multiplex Fluorescent Reagent Kit v2 (ACD, Advanced Cell Diagnostics) was used following the manufacturer’s manual. Probe targeting mouse *Kat7* (#1126701) or *Jade2* (#1725601) was purchased from ACD. Images were collected with a Zeiss LSM 900 confocal microscope and analyzed using ImageJ software (NIH).

### Acute brain slice electrophysiology

6 months old mice were anesthetized with isoflurane, and then perfused intracardially with ice-cold oxygenated cutting solution containing (in mM): 110 choline chloride, 2.5 KCl, 1.25 NaH_2_PO_4_, 25 NaHCO_3_, 0.5 CaCl_2_, 7 MgCl_2_, 10 glucose, saturated with 95% O_2_ / 5% CO_2_. The brain was removed rapidly and immersed in ice-cold oxygenated cutting solution. Transverse hippocampal slices (350 µm) were cut in the cutting solution using a vibratome (VT-1200S, Leica) and transferred to artificial cerebrospinal fluid (aCSF) containing (in mM): 125 NaCl, 2.5 KCl, 1.25 NaH_2_PO_4_, 25 NaHCO_3_, 2 CaCl_2_, 2 MgCl_2_, 10 glucose, saturated with 95% O_2_ / 5% CO_2_. The slices were recovered for 20 min at 35 °C and then maintained at room temperature for 1 hour. Slices were subsequently transferred to a submerged recording chamber containing aCSF solution maintained at 34 °C. Picrotoxin (100 µM) was added to block inhibitory transmission. mEPSCs were recorded at a holding potential of -70 mV in the presence of 1 µM tetrodotoxin (TTX). fEPSPs were evoked in the CA1 stratum radiatum by stimulating the Schaffer collateral with a concentric bipolar electrode and recorded with a glass pipette (1-3 MΩ) filled with aCSF. The stimulus intensity was adjusted to evoke 40%-50% of the maximal response. LTP was induced by theta burst stimulation (TBS) consisting of two trains of 5 bursts at 5 Hz, and each burst contained 4 pulses at 100 Hz. Recordings were made with MultiClamp 700B amplifier (Molecular Devices) and data acquisition was performed with pClamp 10.7 software (Molecular Devices).

### Morris water maze test

Morris water maze tests were performed at Behavioral Core of Johns Hopkins University. In brief, we used a maze consisted of a round pool (diameter, 120 cm) filled with water that was at 24 °C and made opaque with nontoxic white paint. A circular plastic platform (diameter, 10 cm) was placed at the center of the target quadrant and submerged 1 cm below the surface of the water. Four local cues were provided to allow spatial map generation. In brief, we trained the mice for four trials per day with different start points for five consecutive days. Mice were gently placed into the water facing the wall of the pool and allowed to freely explore the whole maze for 1 min. Mice were then guided to the rescue platform if they did not find it. Mice were allowed to take a rest on the platform for 10 s and then retrained from a different start position with the same procedure. The latency for each animal to find the platform (at least 3 s stay) was recorded. On day 6, the platform was removed, and animals searched freely for 1 min starting from the opposite quadrant. The entries into the platform area, total time spent in the target quadrant, and the total distance travels were recorded using the ANY-maze software.

### Statistical analysis

Statistical analyses were performed using GraphPad Prism 9 (GraphPad Software, CA). Before statistical analysis, variation within each group of data and the assumptions of the tests were checked. For in vitro experiments, the cells were evenly suspended and randomly distributed in each well tested. For in vivo experiments, the animals were distributed into various treatment groups randomly. Comparisons between two groups were made using unpaired Student’s two-tailed t test or Mann-Whitney test. Comparisons among three or more groups were made using one- or two-way ANOVA followed by Bonferroni’s post hoc test. The significance level was set at p < 0.05. Test statistics, n numbers, and p values are indicated in the figure legends. All data are presented as mean ± SEM.

## Acknowledgments

We thank Dr. Tony Wyss-Coray for sharing the BV2 cell line, Dr. Rong Wu for kindly providing technical training in immunofluorescence of human patient tissues, Dr. Chaohua Jiang for the technical assistance on recording, and the members of Dr. Qiu lab for valuable discussions.

## Author contributions

Y.L. initiated the project and performed the majority of the experiments. M.F. performed LTP recordings and osmotic pump implantation. Y.Y. and H.Y.C. analyzed the sequencing data. H.Y.C. performed the flow cytometry of isolated microglia. S.S. provided the human brain samples. Y.L., M.F., Y.Y., H.Y.C., and Z.Q. analyzed and interpreted the results. Y.L. and Z.Q. designed the study and wrote the paper with input from all authors.

## REFERENCES

1. Chen, S., Cao, Z., Nandi, A., Counts, N., Jiao, L., Prettner, K., Kuhn, M., Seligman, B., Tortorice, D., Vigo, D., et al. (2024). The global macroeconomic burden of Alzheimer’s disease and other dementias: estimates and projections for 152 countries or territories. Lancet Glob Health 12, e1534–e1543. 10.1016/S2214-109X(24)00264-X.

2. Knopman, D.S., Amieva, H., Petersen, R.C., Chetelat, G., Holtzman, D.M., Hyman, B.T., Nixon, R.A., and Jones, D.T. (2021). Alzheimer disease. Nat Rev Dis Primers 7, 33. 10.1038/s41572-021-00269-y.

3. Self, W.K., and Holtzman, D.M. (2023). Emerging diagnostics and therapeutics for Alzheimer disease. Nat Med 29, 2187–2199. 10.1038/s41591-023-02505-2.

4. Shi, F.D., and Yong, V.W. (2025). Neuroinflammation across neurological diseases. Science 388, eadx0043. 10.1126/science.adx0043.

5. Heneka, M.T., van der Flier, W.M., Jessen, F., Hoozemanns, J., Thal, D.R., Boche, D., Brosseron, F., Teunissen, C., Zetterberg, H., Jacobs, A.H., et al. (2025). Neuroinflammation in Alzheimer disease. Nat Rev Immunol 25, 321–352. 10.1038/s41577-024-01104-7.

6. van der Flier, W.M., and Heneka, M.T. (2025). Zooming in on brain inflammation in Alzheimer’s disease. Brain 148, 1–2. 10.1093/brain/awae394.

7. Karch, C.M., and Goate, A.M. (2015). Alzheimer’s disease risk genes and mechanisms of disease pathogenesis. Biol Psychiatry 77, 43–51. 10.1016/j.biopsych.2014.05.006.

8. Keren-Shaul, H., Spinrad, A., Weiner, A., Matcovitch-Natan, O., Dvir-Szternfeld, R., Ulland, T.K., David, E., Baruch, K., Lara-Astaiso, D., Toth, B., et al. (2017). A Unique Microglia Type Associated with Restricting Development of Alzheimer’s Disease. Cell 169, 1276–1290 e1217. 10.1016/j.cell.2017.05.018.

9. Eskandari-Sedighi, G., and Blurton-Jones, M. (2023). Microglial APOE4: more is less and less is more. Mol Neurodegener 18, 99. 10.1186/s13024-023-00693-6.

10. Leng, F., and Edison, P. (2021). Neuroinflammation and microglial activation in Alzheimer disease: where do we go from here? Nat Rev Neurol 17, 157–172. 10.1038/s41582-020-00435-y.

11. Zhong, F., Liang, S., and Zhong, Z. (2019). Emerging Role of Mitochondrial DNA as a Major Driver of Inflammation and Disease Progression. Trends Immunol 40, 1120–1133. 10.1016/j.it.2019.10.008.

12. Riley, J.S., and Tait, S.W. (2020). Mitochondrial DNA in inflammation and immunity. EMBO Rep 21, e49799. 10.15252/embr.201949799.

13. Marchi, S., Guilbaud, E., Tait, S.W.G., Yamazaki, T., and Galluzzi, L. (2023). Mitochondrial control of inflammation. Nat Rev Immunol 23, 159–173. 10.1038/s41577-022-00760-x.

14. Gulen, M.F., Samson, N., Keller, A., Schwabenland, M., Liu, C., Gluck, S., Thacker, V.V., Favre, L., Mangeat, B., Kroese, L.J., et al. (2023). cGAS-STING drives ageing-related inflammation and neurodegeneration. Nature 620, 374–380. 10.1038/s41586-023-06373-1.

15. Xie, X., Ma, G., Li, X., Zhao, J., Zhao, Z., and Zeng, J. (2023). Activation of innate immune cGAS-STING pathway contributes to Alzheimer’s pathogenesis in 5xFAD mice. Nat Aging 3, 202–212. 10.1038/s43587-022-00337-2.

16. Udeochu, J.C., Amin, S., Huang, Y., Fan, L., Torres, E.R.S., Carling, G.K., Liu, B., McGurran, H., Coronas-Samano, G., Kauwe, G., et al. (2023). Tau activation of microglial cGAS-IFN reduces MEF2C-mediated cognitive resilience. Nat Neurosci 26, 737–750. 10.1038/s41593-023-01315-6.

17. Decout, A., Katz, J.D., Venkatraman, S., and Ablasser, A. (2021). The cGAS-STING pathway as a therapeutic target in inflammatory diseases. Nat Rev Immunol 21, 548–569. 10.1038/s41577-021-00524-z.

18. Chung, S., Jeong, J.H., Park, J.C., Han, J.W., Lee, Y., Kim, J.I., and Mook-Jung, I. (2024). Blockade of STING activation alleviates microglial dysfunction and a broad spectrum of Alzheimer’s disease pathologies. Exp Mol Med 56, 1936–1951. 10.1038/s12276-024-01295-y.

19. Borrelli, E., Nestler, E.J., Allis, C.D., and Sassone-Corsi, P. (2008). Decoding the epigenetic language of neuronal plasticity. Neuron 60, 961–974. 10.1016/j.neuron.2008.10.012.

20. Sweatt, J.D. (2013). The emerging field of neuroepigenetics. Neuron 80, 624–632. 10.1016/j.neuron.2013.10.023.

21. Cavalli, G., and Heard, E. (2019). Advances in epigenetics link genetics to the environment and disease. Nature 571, 489–499. 10.1038/s41586-019-1411-0.

22. Graff, J., and Tsai, L.H. (2013). Histone acetylation: molecular mnemonics on the chromatin. Nat Rev Neurosci 14, 97–111. 10.1038/nrn3427.

23. Yokoyama, A., Niida, H., Kutateladze, T.G., and Cote, J. (2024). HBO1, a MYSTerious KAT and its links to cancer. Biochim Biophys Acta Gene Regul Mech 1867, 195045. 10.1016/j.bbagrm.2024.195045.

24. Pulido-Salgado, M., Vidal-Taboada, J.M., Barriga, G.G., Sola, C., and Saura, J. (2018). RNA-Seq transcriptomic profiling of primary murine microglia treated with LPS or LPS + IFNgamma. Sci Rep 8, 16096. 10.1038/s41598-018-34412-9.

25. Oakley, H., Cole, S.L., Logan, S., Maus, E., Shao, P., Craft, J., Guillozet-Bongaarts, A., Ohno, M., Disterhoft, J., Van Eldik, L., et al. (2006). Intraneuronal beta-amyloid aggregates, neurodegeneration, and neuron loss in transgenic mice with five familial Alzheimer’s disease mutations: potential factors in amyloid plaque formation. J Neurosci 26, 10129–10140. 10.1523/JNEUROSCI.1202-06.2006.

26. Srinivasan, K., Friedman, B.A., Etxeberria, A., Huntley, M.A., van der Brug, M.P., Foreman, O., Paw, J.S., Modrusan, Z., Beach, T.G., Serrano, G.E., and Hansen, D.V. (2020). Alzheimer’s Patient Microglia Exhibit Enhanced Aging and Unique Transcriptional Activation. Cell Rep 31, 107843. 10.1016/j.celrep.2020.107843.

27. MacPherson, L., Anokye, J., Yeung, M.M., Lam, E.Y.N., Chan, Y.C., Weng, C.F., Yeh, P., Knezevic, K., Butler, M.S., Hoegl, A., et al. (2020). HBO1 is required for the maintenance of leukaemia stem cells. Nature 577, 266–270. 10.1038/s41586-019-1835-6.

28. Kueh, A.J., Dixon, M.P., Voss, A.K., and Thomas, T. (2011). HBO1 is required for H3K14 acetylation and normal transcriptional activity during embryonic development. Mol Cell Biol 31, 845–860. 10.1128/MCB.00159-10.

29. Kueh, A.J., Bergamasco, M.I., Quaglieri, A., Phipson, B., Li-Wai-Suen, C.S.N., Lonnstedt, I.M., Hu, Y., Feng, Z.P., Woodruff, C., May, R.E., et al. (2023). Stem cell plasticity, acetylation of H3K14, and de novo gene activation rely on KAT7. Cell Rep 42, 111980. 10.1016/j.celrep.2022.111980.

30. Zhong, Z., Liang, S., Sanchez-Lopez, E., He, F., Shalapour, S., Lin, X.J., Wong, J., Ding, S., Seki, E., Schnabl, B., et al. (2018). New mitochondrial DNA synthesis enables NLRP3 inflammasome activation. Nature 560, 198–203. 10.1038/s41586-018-0372-z.

31. Xian, H., and Karin, M. (2023). Oxidized mitochondrial DNA: a protective signal gone awry. Trends Immunol 44, 188–200. 10.1016/j.it.2023.01.006.

32. Xian, H., Watari, K., Sanchez-Lopez, E., Offenberger, J., Onyuru, J., Sampath, H., Ying, W., Hoffman, H.M., Shadel, G.S., and Karin, M. (2022). Oxidized DNA fragments exit mitochondria via mPTP- and VDAC-dependent channels to activate NLRP3 inflammasome and interferon signaling. Immunity 55, 1370–1385 e1378. 10.1016/j.immuni.2022.06.007.

33. Miura, H., Quadros, R.M., Gurumurthy, C.B., and Ohtsuka, M. (2018). Easi-CRISPR for creating knock-in and conditional knockout mouse models using long ssDNA donors. Nat Protoc 13, 195–215. 10.1038/nprot.2017.153.

34. Sahasrabuddhe, V., and Ghosh, H.S. (2022). Cx3Cr1-Cre induction leads to microglial activation and IFN-1 signaling caused by DNA damage in early postnatal brain. Cell Rep 38, 110252. 10.1016/j.celrep.2021.110252.

35. Yum, S., Li, M., Fang, Y., and Chen, Z.J. (2021). TBK1 recruitment to STING activates both IRF3 and NF-kappaB that mediate immune defense against tumors and viral infections. Proc Natl Acad Sci U S A 118. 10.1073/pnas.2100225118.

36. Venegas, C., Kumar, S., Franklin, B.S., Dierkes, T., Brinkschulte, R., Tejera, D., Vieira-Saecker, A., Schwartz, S., Santarelli, F., Kummer, M.P., et al. (2017). Microglia-derived ASC specks cross-seed amyloid-beta in Alzheimer’s disease. Nature 552, 355–361. 10.1038/nature25158.

37. Hur, J.Y., Frost, G.R., Wu, X., Crump, C., Pan, S.J., Wong, E., Barros, M., Li, T., Nie, P., Zhai, Y., et al. (2020). The innate immunity protein IFITM3 modulates gamma-secretase in Alzheimer’s disease. Nature 586, 735–740. 10.1038/s41586-020-2681-2.

38. Tsien, J.Z., Chen, D.F., Gerber, D., Tom, C., Mercer, E.H., Anderson, D.J., Mayford, M., Kandel, E.R., and Tonegawa, S. (1996). Subregion- and cell type-restricted gene knockout in mouse brain. Cell 87, 1317–1326. 10.1016/s0092-8674(00)81826-7.

39. Orihuela, R., McPherson, C.A., and Harry, G.J. (2016). Microglial M1/M2 polarization and metabolic states. Br J Pharmacol 173, 649–665. 10.1111/bph.13139.

40. Sangineto, M., Ciarnelli, M., Cassano, T., Radesco, A., Moola, A., Bukke, V.N., Romano, A., Villani, R., Kanwal, H., Capitanio, N., et al. (2023). Metabolic reprogramming in inflammatory microglia indicates a potential way of targeting inflammation in Alzheimer’s disease. Redox Biol 66, 102846. 10.1016/j.redox.2023.102846.

41. Natarajan, N., Florentin, J., Johny, E., Xiao, H., O’Neil, S.P., Lei, L., Shen, J., Ohayon, L., Johnson, A.R., Rao, K., et al. (2024). Aberrant mitochondrial DNA synthesis in macrophages exacerbates inflammation and atherosclerosis. Nat Commun 15, 7337. 10.1038/s41467-024-51780-1.

42. Guan, X., Zhu, S., Song, J., Liu, K., Liu, M., Xie, L., Wang, Y., Wu, J., Xu, X., and Pang, T. (2024). Microglial CMPK2 promotes neuroinflammation and brain injury after ischemic stroke. Cell Rep Med 5, 101522. 10.1016/j.xcrm.2024.101522.

43. Guan, J.S., Haggarty, S.J., Giacometti, E., Dannenberg, J.H., Joseph, N., Gao, J., Nieland, T.J., Zhou, Y., Wang, X., Mazitschek, R., et al. (2009). HDAC2 negatively regulates memory formation and synaptic plasticity. Nature 459, 55–60. 10.1038/nature07925.

44. Mews, P., Donahue, G., Drake, A.M., Luczak, V., Abel, T., and Berger, S.L. (2017). Acetyl-CoA synthetase regulates histone acetylation and hippocampal memory. Nature 546, 381–386. 10.1038/nature22405.

45. Liu, Y., Fan, M., Yang, J., Mihaljevic, L., Chen, K.H., Ye, Y., Sun, S., and Qiu, Z. (2024). KAT6A deficiency impairs cognitive functions through suppressing RSPO2/Wnt signaling in hippocampal CA3. Sci Adv 10, eadm9326. 10.1126/sciadv.adm9326.

46. Zhang, K., Schrag, M., Crofton, A., Trivedi, R., Vinters, H., and Kirsch, W. (2012). Targeted proteomics for quantification of histone acetylation in Alzheimer’s disease. Proteomics 12, 1261–1268. 10.1002/pmic.201200010.

47. Plagg, B., Ehrlich, D., Kniewallner, K.M., Marksteiner, J., and Humpel, C. (2015). Increased Acetylation of Histone H4 at Lysine 12 (H4K12) in Monocytes of Transgenic Alzheimer’s Mice and in Human Patients. Curr Alzheimer Res 12, 752–760. 10.2174/1567205012666150710114256.

48. Nativio, R., Donahue, G., Berson, A., Lan, Y., Amlie-Wolf, A., Tuzer, F., Toledo, J.B., Gosai, S.J., Gregory, B.D., Torres, C., et al. (2018). Dysregulation of the epigenetic landscape of normal aging in Alzheimer’s disease. Nat Neurosci 21, 497–505. 10.1038/s41593-018-0101-9.

49. Wang, W., Zheng, Y., Sun, S., Li, W., Song, M., Ji, Q., Wu, Z., Liu, Z., Fan, Y., Liu, F., et al. (2021). A genome-wide CRISPR-based screen identifies KAT7 as a driver of cellular senescence. Sci Transl Med 13. 10.1126/scitranslmed.abd2655.

50. Xiao, Y., Li, W., Yang, H., Pan, L., Zhang, L., Lu, L., Chen, J., Wei, W., Ye, J., Li, J., et al. (2021). HBO1 is a versatile histone acyltransferase critical for promoter histone acylations. Nucleic Acids Res 49, 8037–8059. 10.1093/nar/gkab607.

51. Zhou, Y., Jia, K., Wang, S., Li, Z., Li, Y., Lu, S., Yang, Y., Zhang, L., Wang, M., Dong, Y., et al. (2023). Malignant progression of liver cancer progenitors requires lysine acetyltransferase 7-acetylated and cytoplasm-translocated G protein GalphaS. Hepatology 77, 1106–1121. 10.1002/hep.32487.

52. Marschallinger, J., Iram, T., Zardeneta, M., Lee, S.E., Lehallier, B., Haney, M.S., Pluvinage, J.V., Mathur, V., Hahn, O., Morgens, D.W., et al. (2020). Lipid-droplet-accumulating microglia represent a dysfunctional and proinflammatory state in the aging brain. Nat Neurosci 23, 194–208. 10.1038/s41593-019-0566-1.

53. Liu, Y., Lai, S., Ma, W., Ke, W., Zhang, C., Liu, S., Zhang, Y., Pei, F., Li, S., Yi, M., et al. (2017). CDYL suppresses epileptogenesis in mice through repression of axonal Nav1.6 sodium channel expression. Nat Commun 8, 355. 10.1038/s41467-017-00368-z.

