## Supplemental Figure 1-8 for "Epigenetic control of microglial mitochondrial immunity by KAT7 drives Alzheimer’s disease pathogenesis"

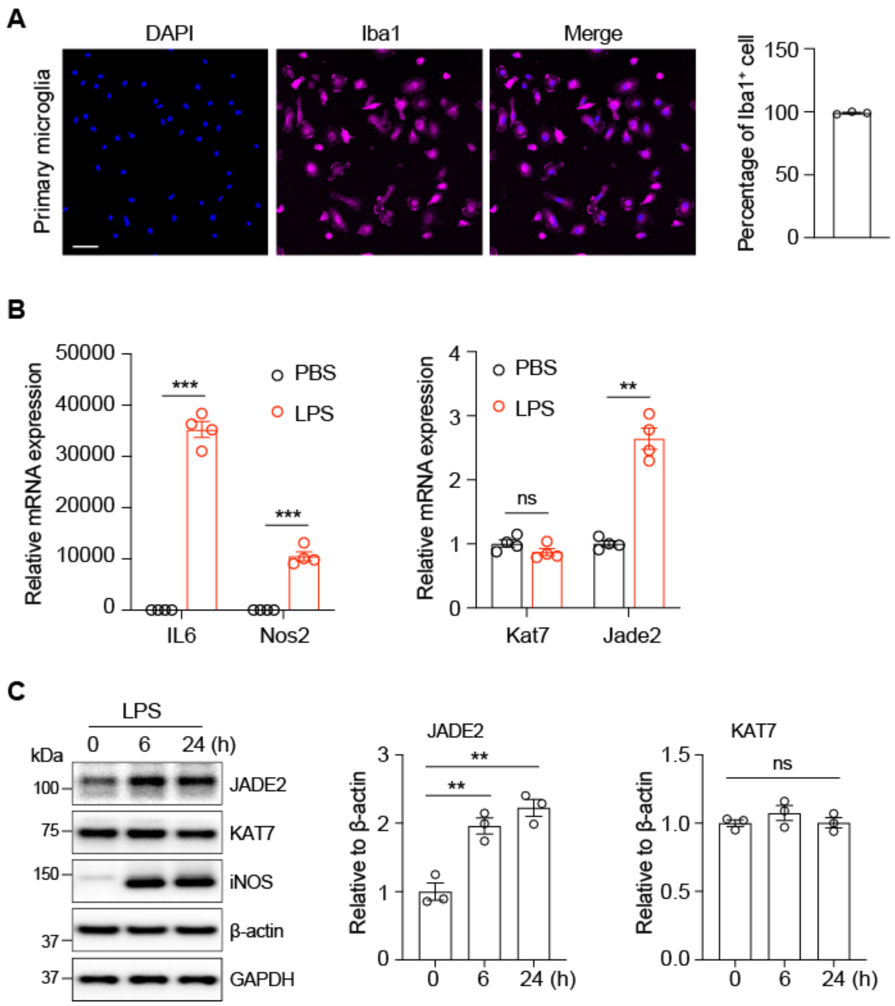


**Suppl. Fig. 1. The expression of *Jade2* is upregulated in LPS-induced microglial activation**. **A**, Representative images (left) and quantification (right) of Iba1 staining in cultured mouse primary microglia. Scale bar, 40 µm. n=3. **B**, qPCR analysis of indicated genes in primary microglia treated with LPS for 6 h. n=4, two-way ANOVA test. **C**, Western blot analysis of indicated protein in BV2 cells treated with LPS. Quantification was done by normalizing to β-actin (right). n=3, one-way ANOVA test. **p<0.01, ***p<0.001. Data are mean±SEM.

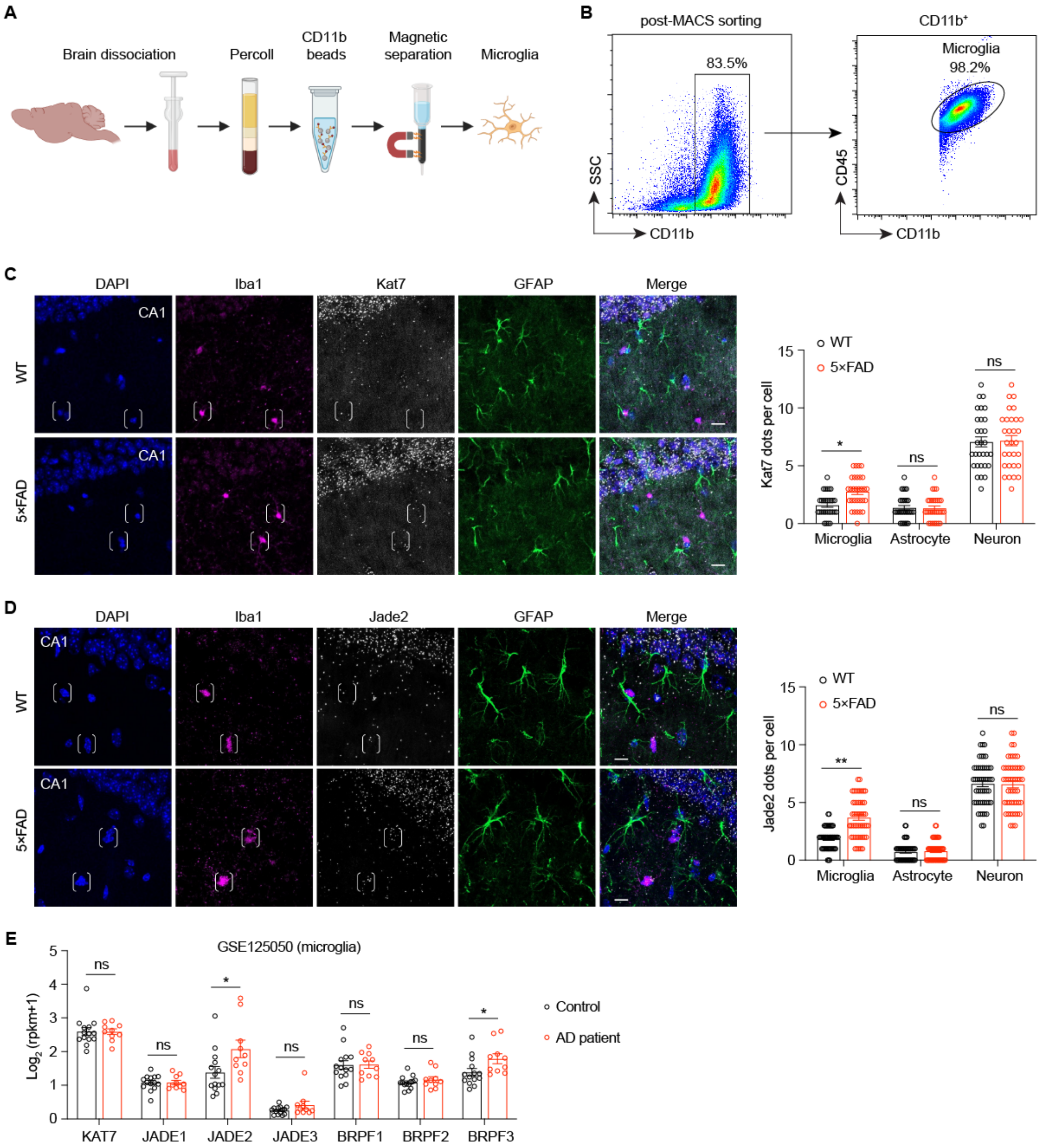


**Suppl. Fig. 2. Expression of the KAT7 complex is elevated in microglia from both 5×FAD mouse model and human AD patients**. **A**, Schematic diagram of microglia isolated from adult mouse brains. **B**, Representative FACS plots and gating strategy to check the purity of CD11b isolated microglia from the adult mouse brain. **C-D**, Left: Representative images of *Kat7* (**C**) or *Jade2* (**D**) RNAscope and its colocalization with Iba1- and GFAP-positive cells in hippocampal CA1 region of 7-month-old WT and 5×FAD mice. Scale bar, 10 μm. Right: Quantification in different cell types. n=30-50 cells from 3 mice per group. Two-way ANOVA test. **E**, Upregulation of JADE2 and BRPF3 in microglia from AD patients based on GSE125050 dataset. n= 14 control and 10 AD. Unpaired student's *t*-test. *$p<0.05$, **$p<0.01$. ns, nonsignificant. Data are mean±SEM.

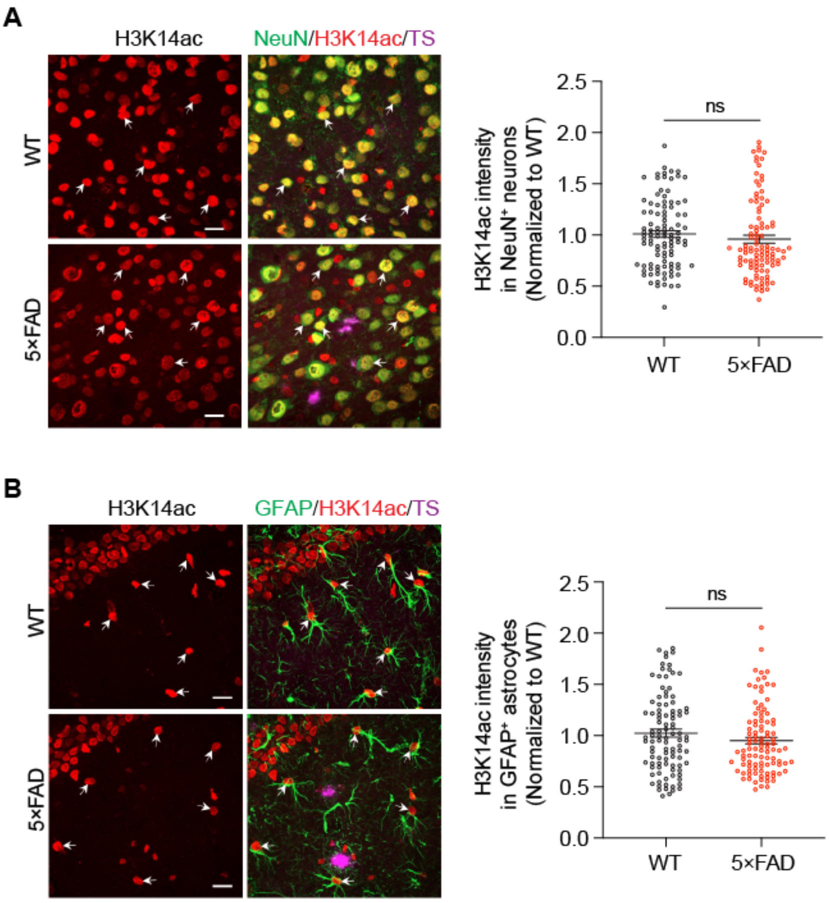


**Suppl. Fig. 3. H3K14ac levels are not changed in neurons or astrocytes of 5×FAD mice.** **A**, Left: Representative images of H3K14ac co-stained with neurons (NeuN) in the cortex region of 6-month-old WT and 5×FAD mice. Scale bar, 20 μm. Right: Quantification of H3K14ac intensity in neurons (n=100 cells from 3 mice per group). White arrowheads indicate H3K14ac in neurons. Mann-Whitney test. **B**, Left: Representative images of H3K14ac co-stained with astrocytes (GFAP) in the cortex region of 6-month-old WT and 5×FAD mice. Scale bar, 20 μm. Right: Quantification of H3K14ac intensity in astrocytes (n=100 cells from 3 mice per group). White arrowheads indicate H3K14ac in astrocytes. Mann-Whitney test. ns, nonsignificant. Data are mean±SEM.

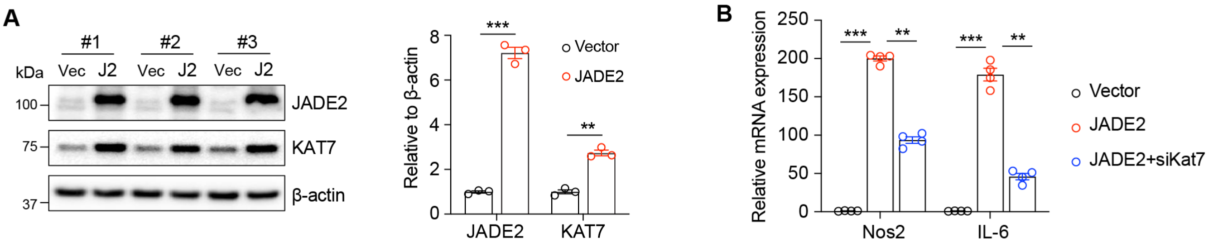


**Suppl. Fig. 4. JADE2 overexpression enhances expression of pro-inflammatory factors.** **A**, Western blot analysis of JADE2 overexpression in BV2 cells. Quantification was done by normalizing to β-actin (right). n=3. **B**, qPCR analysis of *Nos2* and *Il-6* levels in BV2 cells. n=4. **p<0.01, ***p<0.001. Two-way ANOVA test. Data are mean±SEM.

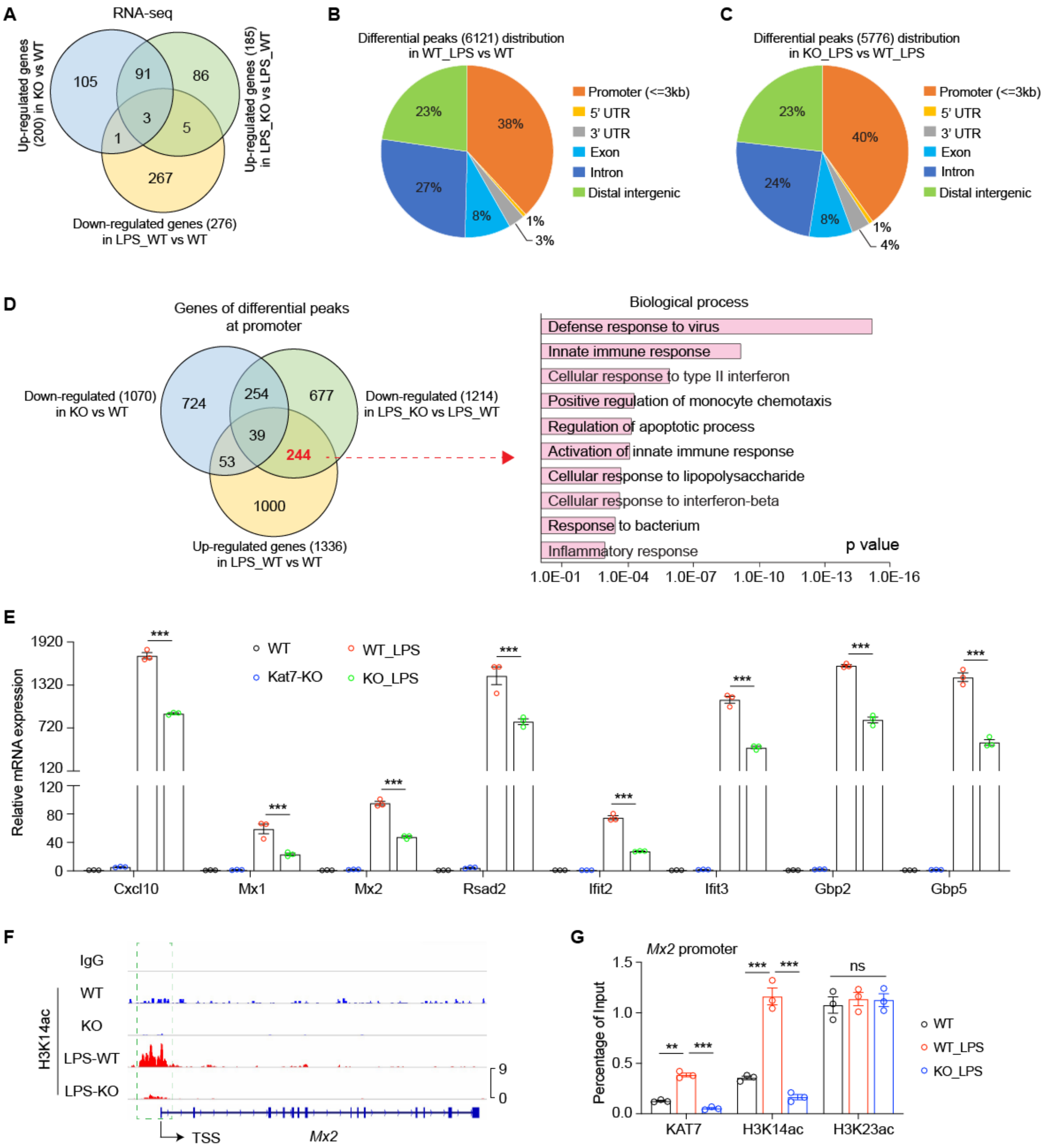


**Suppl. Fig. 5. Analysis and validation of RNA-seq and CUT&Tag data**. **A**, Venn diagram of overlapping genes in RNA-seq. **B**, Genomic distribution of differential H3K14ac-binding peaks in WT_LPS vs WT. **C**, Genomic distribution of differential H3K14ac-binding peaks in KO_LPS vs WT_LPS. **D**, Left: Venn diagram of overlapped genes of differential peaks at promoter among downregulated in KO vs WT, downregulated in KO_LPS vs WT_LPS, and upregulated in WT_LPS vs WT. Right: GO pathway analysis of the 244 overlapped genes. **E**, qPCR analysis showed the expression of the indicated genes in BV2 microglia with or without LPS treatment. n=3. **F**, Representative CUT&Tag tracks of H3K14ac in *Mx2*. Green box indicated proximal promoter. TSS, transcriptional start site. **G**, qChIP analysis of *Mx2* promoter using the indicated antibodies in BV2 cells. n=3. **p<0.01, ***p<0.001. Two-way ANOVA test. Data are mean±SEM.

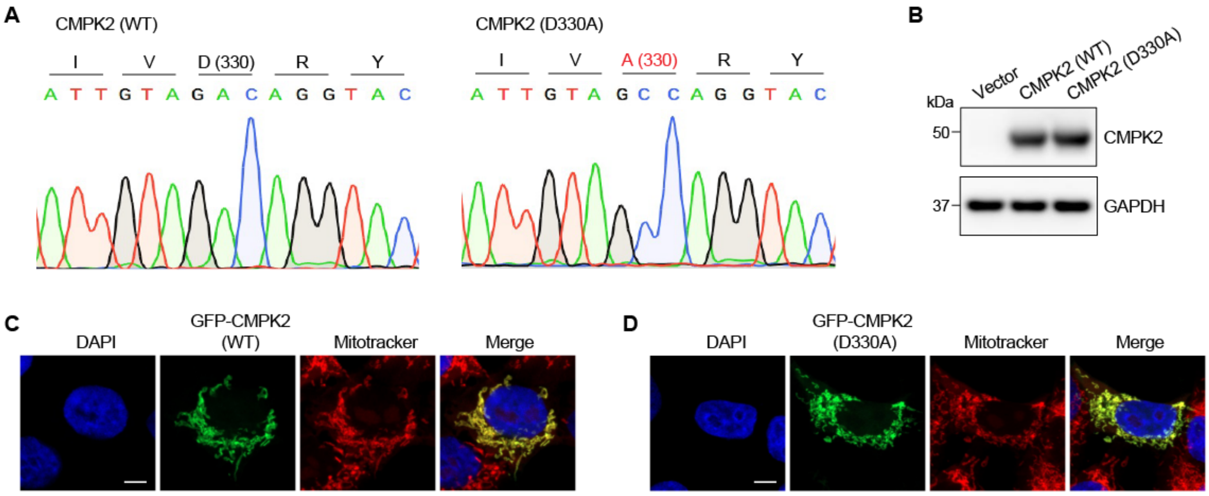


**Suppl. Fig. 6. The CMPK2 mutant is still expressed in mitochondria. A**, Sanger sequencing validated the CMPK2 mutant (D330A) sequence. **B**, Western blot confirmed the infection efficiency of CMPK2 WT and mutant lentiviruses in BV2 microglia. **C-D**, Representative images of GFP-CMPK2 (**C**) or GFP-CMPK2-D330A (**D**) co-stained with Mitotracker in HEK-293T cells.

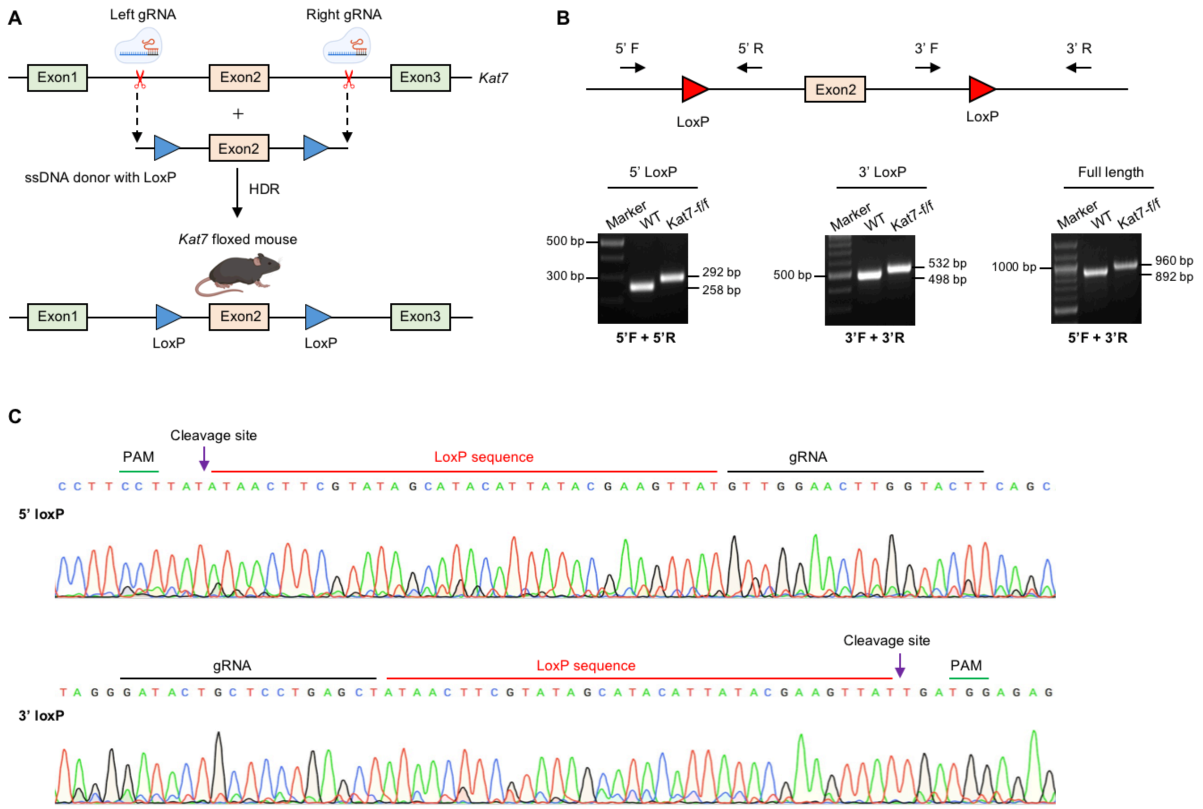


**Suppl. Fig. 7. Generation of *Kat7*-floxed mice**. **A**, Scheme of *Kat7*-floxed mice generation using CRISPR-Cas9 method. HDR, homology-directed repair. **B-C**, Validation of *Kat7*-floxed mice by mouse tail genotyping (**B**) and Sanger sequencing (**C**).

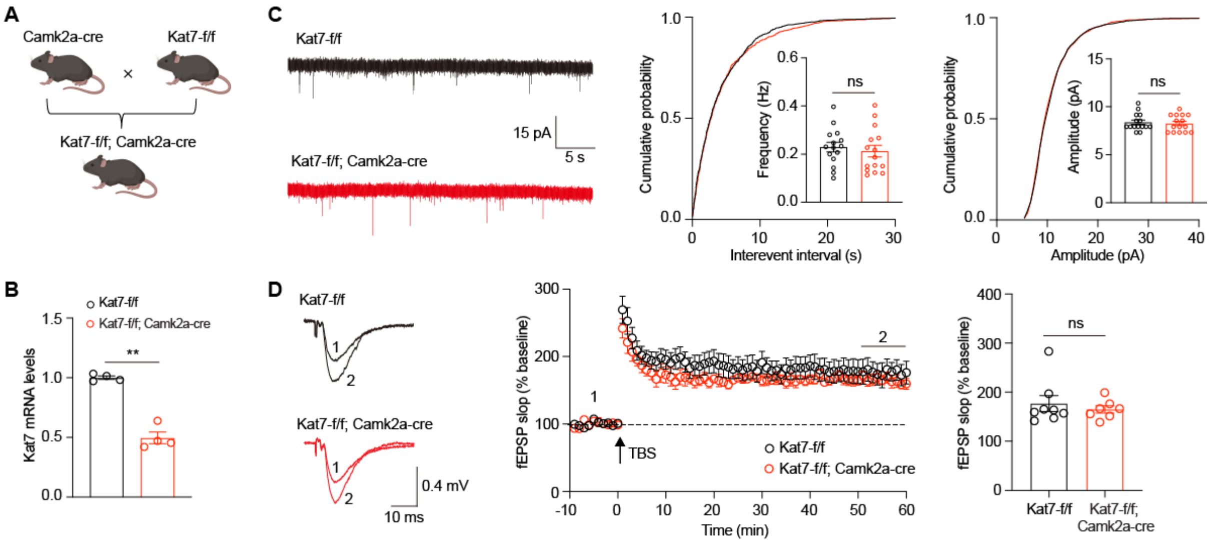


**Suppl. Fig. 8. Deletion of *Kat7* in excitatory neurons does not alter synaptic function in mice**. **A**, *Kat7* floxed mice crossed with Camk2a-cre mice to obtain the $Kat7^{\text{Camk2a-cKO}}$ mice. **B**, qPCR analysis of total hippocampal mRNA from $Kat7^{\text{f/f}}$ and $Kat7^{\text{Camk2a-cKO}}$ mice. n=4 mice per group. **C**, Representative traces (left) and quantification of frequency (middle) and amplitude (right) of mEPSC in hippocampal CA1 pyramidal neurons. n=15 cells from 3 mice per group. **D**, TBS-induced LTP at Schaffer collateral to CA1 synapses. Arrow indicates LTP induction. n=7-8 slices from 4 mice per group. Mann-Whitney test. ns, nonsignificant. Data are mean± SEM.

**Table S1**. The information for formalin-fixed paraffin embedded (FFPE) tissue slides of frontal cortex from control and AD patients.

| Case | Patient GUID | Age | Gender | Tissue |
|---|---|---|---|---|
| Control 1 | BRC2052 | 79 | Male | Middle frontal gyrus |
| Control 2 | BRC2151 | 72 | Male | Middle frontal gyrus |
| Control 3 | BRC2234 | 68 | Female | Middle frontal gyrus |
| Control 4 | BRC2317 | 65 | Female | Middle frontal gyrus |
| AD 1 | BRC2609 | 68 | Male | Middle frontal gyrus |
| AD 2 | BRC2636 | 80 | Female | Middle frontal gyrus |
| AD 3 | BRC2718 | 70 | Male | Middle frontal gyrus |
| AD 4 | BRC2641 | 72 | Female | Middle frontal gyrus |

**Table S2**. Mouse genotyping primers

| Mouse strain | Strand | Sequence |
|---|---|---|
| Kat7-floxed mice | 5’F | AGTACAGGTGGTTTGGTTGT |
| | 5’R | AATCGGAATCTTCGGTTCCA |
| | 3’F | CACAGTGGACAGTGGTGTCA |
| | 3’R | TCAGCAGCTGCCTTACACTT |
| Cx3cr1-$Cre^{ER}$ | F (common) | AAGACTCACGTGGACCTGCT |
| | R (WT) | AGGATGTTGACTTCCGAGTTG |
| | R ($Cre^{ER}$) | CGGTTATTCAACTTGCACCA |
| Camk2a-Cre | F | GTTCTCCGTTTGCACTCAGG |
| | R | CAGGTTCTTGCGAACCTCAT |
| 5×FAD | F (common) | ACCCCCATGTCAGAGTTCCT |
| | R (WT) | TATACAACCTTGGGGGATGG |
| | R (mutant) | CGGGCCTCTTCGCTATTAC |

**Table S3**. Real-time qPCR primers

| Mouse gene name | Strand | Sequence |
|---|---|---|
| *Kat7* | F | TGCAGGCAGTAGTTCAGATGG |
| | R | CAGGGCTGGAATCTTGGGAA |
| *Jade1* | F | ATGTCTGCCAGTCACCTGATGG |
| | R | ACGACATAGCCAACTGCCCTCT |
| *Jade2* | F | CCAAGACTGACGAGGTGGACAA |
| | R | CCGTCTTGTCACCATGTAGCAC |
| *Jade3* | F | GTCTCCAGACAGTGAAGAAGGG |
| | R | GCACAGCCAACTACCTTCTGGA |
| *Brpf1* | F | CCAAGAGAAGGACACAGGCAAC |
| | R | GTAGCGGTAAGCCTCCAAGTTC |
| *Brpf2* | F | GGAGGCTTTGAAGATGAGGCTG |
| | R | GGTGGTTCTGAGTTAGTCTCCG |

| | | |
|---|---|---|
| *Brpf3* | F | CTCATCCGCAAAAGGGAGAAGC |
| | R | TCCAGAGTCGTCCTCAACAGGA |
| *Nos2* | F | CAGCTGGGCTGTACAAACCTT |
| | R | CATTGGAAGTGAAGCGTTTCG |
| *Il-6* | F | CTGCAAGAGACTTCCATCCAG |
| | R | AGTGGTATAGACAGGTCTGTTGG |
| *Cmpk2* | F | AACTCTGCGGTGTTCCAAGACC |
| | R | GGAACTTCCCTTTCTGGACCTC |
| *Cxcl10* | F | ATCATCCCTGCGAGCCTATCCT |
| | R | GACCTTTTTTGGCTAAACGCTTTC |
| *Rsad2* | F | GGAAGGTTTTCCAGTGCCTCCT |
| | R | ACAGGACACCTCTTTGTGACGC |
| *Ifit2* | F | CGAACTACCGTCTGGATGACTG |
| | R | CTTCAACCAGCGCCATTGCTTG |
| *Ifit3* | F | GCTCAGGCTTACGTTGACAAGG |
| | R | CTTTAGGCGTGTCCATCCTTCC |
| *Gbp2* | F | AGATGCCCACAGAAACCCTCCA |
| | R | AAGGCATCTCGCTTGGCTACCA |
| *Gbp5* | F | GAACGCCAAAGAAACAGTGAGCC |
| | R | CTTCCTGGATGCGAATAGCCTC |
| *Mx1* | F | TGGACATTGCTACCACAGAGGC |
| | R | TTGCCTTCAGCACCTCTGTCCA |
| *Mx2* | F | ACCAGAGTGCAAGTGAGGAGCT |
| | R | GTACTAGGGCAGTGATGTCCTG |
| *β-actin* | F | GGCTGTATTCCCCTCCATCG |
| | R | CCAGTTGGTAACAATGCCATGT |
| *Gapdh* | F | GGGTGTGAACCACGAGAAATA |
| | R | CTGTGGTCATGAGCCCTTC |

**Table S4**. mtDNA qPCR primers

| **Gene name** | **Strand** | **Sequence** |
|---|---|---|
| Mouse *D-loop* | F | AATCTACCATCCTCCGTGAAACC |
| | R | TCAGTTTAGCTACCCCCAAGTTTAA |
| Mouse *Nd4* | F | AACGGATCCACAGCCGTA |
| | R | AGTCCTCGGGCCATGATT |
| Mouse *B2m* | F | ATGGGAAGCCGAACATACTG |
| | R | CAGTCTCAGTGGG GGTGAAT |
| Mouse *Tert* | F | TAGCTCATGTGTCAAGACCCTCTT |
| | R | GCCAGCACGTTTCTCTCGTT |

**Table S5**. ChIP-qPCR primers

| Gene name | Strand | Sequence |
|---|---|---|
| Mouse *Cmpk2* promoter | F | ATCCCACGTGAGCAAAGGG |
| | R | CCTGAAGCGTATTTGGGCAG |
| Mouse *Mx2* promoter | F | AGTTCCCCAAGAACCAGAGAAATG |
| | R | ATTATGATGGGAAAGGCAGGTTCA |
